# The N-terminus of YY1 regulates DNA and RNA binding affinity for both the zinc-fingers and an unexpected nucleic acid binding domain

**DOI:** 10.1101/2024.10.04.616721

**Authors:** Jimmy Elias, Jane J. Rosin, Amanda J. Keplinger, Alexander J. Ruthenburg

## Abstract

Transcription factors (TFs) play central roles in dictating cellular identity and function by regulating gene expression programs. Beyond their well-folded DNA binding domains (DBDs) which recognize cognate DNA elements in the genome, TFs are enriched for intrinsically disordered regions (IDRs), which have a host of proposed functions including facilitating protein-protein interactions, aiding in binding site search, and binding RNA. Defining intrinsic regulatory properties of TFs requires further mechanistic investigation. We chose to investigate the DNA and RNA binding properties of Yin Yang 1 (YY1), a ubiquitously expressed TF directly involved in transcriptional activation, repression and genome architecture. Through systematic in vitro nucleic acid binding experiments we resolve conflicting literature defining the RNA binding interface of YY1, demonstrating that there are two RNA binding domains within YY1: its canonical 4 zinc finger DBD and a previously unannotated nucleic acid binding domain, which we term the REPO-NAB. Furthermore, we discover surprising autoinhibitory properties that the N-terminus of the protein imparts on each of these binding domains. Our results provide a new example of IDR-mediated regulation within TFs and enables future mechanistically precise functional investigations.

## INTRODUCTION

Transcription factors (TFs) play a central role in dictating cellular identity (1, 2). Typically modular in composition, TFs are minimally composed of a DNA-binding domain, which imparts sequence-specific DNA recognition, and an activation/effector domain, which interfaces with nuclear protein complexes to recruit or stabilize their local activity (3, 4). Yet for a given TF, only a fraction of accessible cognate sites in the genome are occupied (3, 5–7). Precisely how TFs are localized to subsets of regulatory genetic elements distributed throughout the genome to define distinct gene expression programs remains unclear (7–10). The prevailing explanation for this disconnect is that there are additional genomic interfaces, either through DNA, RNA, or co-factor proteins that further specify localization through multivalent energetics (3, 4, 11–15). Yet for most transcription factors, detailed mapping of these interfaces and their specificities is lacking such that the conventional explanation for site-specificity remains untested. To begin to explicitly test these ideas, we chose to biochemically dissect the nucleic acid binding interface(s) of the constitutively expressed transcription factor Yin Yang 1 (YY1) (16–18), and define the intrinsic autoregulatory features within the protein itself. YY1 derives its name from initial observations describing the TF’s ability to be both a transcriptional activator and repressor of the adeno-associated virus P5 (AAVP5) element (16). While this distinction is attributed to the presence or absence of the E1A cofactor, how YY1 accomplishes these diametrically opposed transcriptional activities in other cases has been a matter of long-standing interest. Context-dependent mechanisms ranging from the ability to interact with core transcriptional machinery (19), engage in DNA repair (20), recruit histone modifying complexes (21–23), and promote through-space contact of cell-type specific enhancers and promoters (18) have all been attributed to this multifunctional TF.

The canonical DNA binding domain of YY1 consists of 4 C_2_H_2_ zinc fingers (ZnFs) (24) situated at the C-terminal end of the protein (25), while the N-terminal portion harbours regions of the protein mapped to activation and/or repressive activities annotated by specific functional studies (16, 17, 19, 21–23) (Fig 1A). In addition to DNA binding, YY1 is capable of binding RNA (26–32). However, there is a standing controversy within the literature regarding the RNA binding interface of YY1 (26–28) rendering experiments that precisely perturb this interface to ascertain the regulatory impact fraught. Previous work has postulated that RNA binding plays a role in both YY1’s homo-dimerization (18), as well as mediating a transcriptional positive feedback loop, where transcriptional machinery is maintained in the local vicinity of gene regulatory elements upon proper transcriptional firing (27). Unambiguous delineation of the impact that RNA binding can have upon YY1 cellular function requires systematic side-by-side comparisons of nucleic acid affinities of putative RNA binding domains compared to full length YY1, and such analyses have not yet been performed.

**Figure 1.**
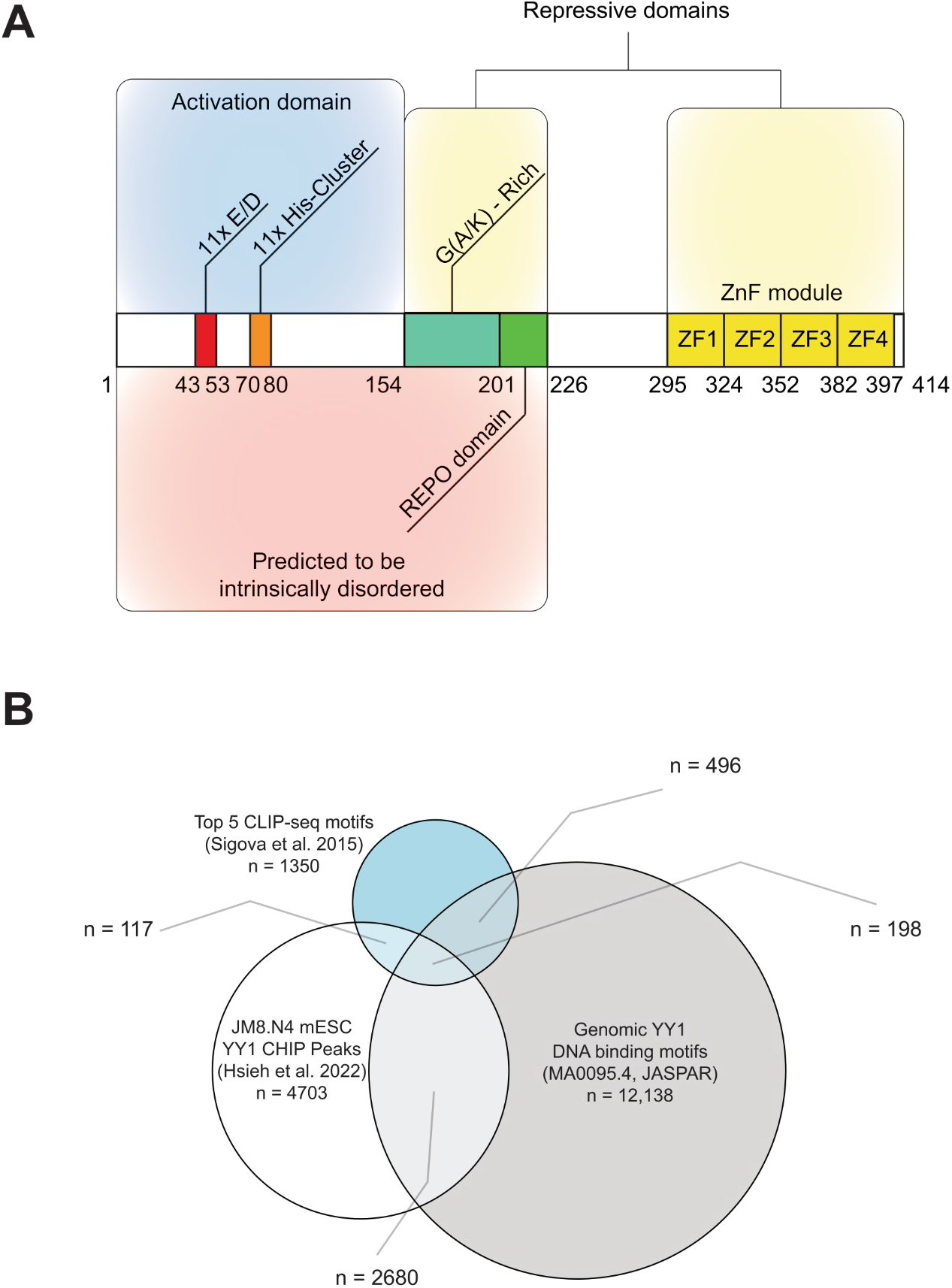
YY1 is a multi-functional transcription factor that adheres to the transcription factor occupancy paradox. **(A)** Schematic of YY1, highlighting domains and key functional features. **(B)** Venn diagram showing the overlap of murine YY1 ChIP-seq sites (34), cognate DNA binding sites (JASPAR, MA0095.4), and RNA sequence motifs (27), all within ATAC accessible loci (33).

Although YY1 is ubiquitously expressed across human cell types and consistently occupies cell-type specific enhancers and promoters, the dominant factors which play a role in determining YY1 genomic occupancy have remained elusive. By taking a nucleic acid-centric perspective and evaluating YY1 binding events from multiple data sets (27, 33, 34), we observe that the genomic occupancy of YY1 cannot be explained by a combination of its cognate dsDNA binding motifs defined by SELEX nor its possible RNA binding motifs derived from CLIP-seq (Fig 1B). These observations prompted our investigation into previously unannotated characteristics of the TF.

In this study, we measure the nucleic acid binding capacity of purified YY1 and fragments thereof using fluorescence polarization. We observe unexpected binding properties for multiple domains of YY1: (1) YY1’s canonical DNA binding ZnF domain exhibits a nearly 10-fold higher affinity for single stranded RNA than double stranded DNA consensus motifs and both nucleic acid types share an overlapping interface; (2) YY1’s Recruitment of Polycomb (REPO) domain (21) possesses previously unannotated tight binding capacity for both DNA and RNA with little apparent sequence specificity. (3) The N-terminus of YY1, which is predicted to be disordered, can inhibit nucleic acid binding to the ZnFs and REPO domain, suggesting an autoinhibitory intra/inter-molecular mechanism to provide fine-tuning of the protein’s activity, and this property seems to account for the weaker apparent affinity for RNA displayed by the full length YY1 protein than displayed for the individual nucleic acid binding domains. Our study elucidates a previously unannotated nucleic acid binding domain of YY1, defines several surprising autoregulatory features intrinsic to the protein which will enable functional tests of their properties *in vivo*, and provides a template for further mechanistic dissection of transcription factors.

## MATERIAL AND METHODS

### Cloning and purification of YY1 constructs

YY1 protein was purified using a method modified from (26). Briefly, a plasmid containing full length YY1 was a gift from Richard Young (Addgene plasmid # 104396; http://n2t.net/addgene:104396; RRID:Addgene_104396) and the full length protein sequence was inserted into a modified pGEX-6P vector using BamHI/XhoI restriction enzyme sites. All proteins were expressed as R3C-cleavable N-terminal fusions with glutathione-S-transferase (GST). Truncation mutants were generated using QuickChange and HiFi methodologies according to manufacturer’s guidelines, with oligonucleotides described in Supplemental Table 1. Protein expression constructs were transformed into Rosetta 2 (DE3) pLysS competent cells and grown in 1L LB cultures containing 34 µg/mL chloramphenicol and 100 µg/mL carbenicillin at 37°C until an optical density at 600 nm (OD_600nm_) measured ∼ 0.6. Upon reaching this OD_600nm_ cultures were cooled to 25°C then induced with 1 mM isopropyl-β-D thiogalactopyranoside (IPTG) and supplemented to 50 µM ZnSO_4_, then shaken at 25°C for 18 hours. Cultures were then harvested by centrifugation in a Thermo Sorvall RC 3BP+ at 4000g for 25 min at 4°C, and resuspended in 50 mLs of Buffer A (20 mM Tris·HCL pH = 8.0, 100 mM NaCl, 5% glycerol) supplemented with 5 mM β-mercaptoethanol, 2 mM dithiothreitol (DTT), 1 mM phenylmethylsulfonyl fluoride (PMSF), and 1250 U of Benzonase (Millipore, 71205-3). Lysis was accomplished via 4 passages through an American Laboratories French Pressure cell, then clarified by centrifugation on a Sorvall 3B+ at 25,000g for 25 min at 4°C. All proteins were purified to homogeneity by the sequence of Glutathione Sepharose 4FF (GE Healthcare, 25 mL bed in XJ-50 column) and Heparin HiTrap (GE Healthcare, 5 mL) chromatography, followed by anion/cation exchange chromatography dependent on the isoelectric point of the respective construct. All protein purification workflows are depicted in Figure S1. HRV-3C protease was utilized to cleave the GST tag after the initial GST column. The size and purity of purified proteins were monitored by SDS-PAGE and protein concentrations were determined by the RC DC Protein Assay (Bio-Rad, 5000122).

### Fluorescence polarization

We used 5’ FAM labeled oligonucleotides as the binding substrates in our fluorescence polarization (FP) assays (sequences are listed in Supplemental Table 1). The RNA and DNA oligonucleotides were resuspended in minimal YY1 binding buffer (10 mM HEPES·KOH [pH = 7.3], 50 mM NaCl, 50 mM KCl, 5 mM MgCl_2_). Duplex DNA substrates were annealed by heating 1.1:1 molar ratio of unlabeled:fluorescently labeled ssDNA oligonucleotides in 1x NATE buffer (50 mM NaCl, 10 mM Tris·HCl [pH = 7.5], 1 mM EDTA) to 95°C followed by slow cooling to room temperature. Purified proteins were dialyzed into YY1 Binding Buffer (10 mM HEPES·KOH [pH = 7.3], 50 mM NaCl, 50 mM KCl, 5 mM MgCl_2,_ 100 µM ZnSO_4_, 0.01% NP-40), quantified, and nucleic acid substrates were added, to a final concentration of either 10 or 20 nM, to the highest protein concentration within the experiment. This protein-nucleic acid solution was then serially diluted with YY1 binding buffer containing the same concentration of nucleic acid, to a final protein concentration of 10 nM. Concentration ranges are stated within the figure legends for each binding experiment. The reaction volumes for each serial dilution were 165 µL, and this reaction was split into three 50 µL technical replicates within black, nonbinding surface 384 well plates (Corning CLS3575). Fluorescence anisotropy was measured by a TECAN Infinite 200 Pro using excitation/emission wavelengths of 485 nm/535 nm with excitation/emission bandwidths of 25 nm/35 nm at 25°C. YY1 binding buffer was used as a blank and G-factor calibration was performed on YY1 binding buffer containing the proper concentration of labeled nucleic acid to produce a fluorescence polarization reading of 20 mP.

Anisotropy data was then analysed in R and fit to nonlinear regressions derived from the Langmuir equation 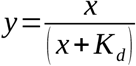 and the Hill equation 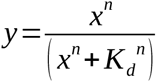 where y is the anisotropy measurement, x is the concentration of purified protein, and the parameters we are fitting for are *K*_*d*_, the dissociation constant, and n, the Hill coefficient. The error in the fit is reported as uncertainty and all reported *K*_*d*_ values have correlation coefficients ≥ 0.9. We chose to present the Langmuir *K*_*d*_ values as the binding is anticipated to be monovalent based on substrate design.

### Circular dichroism

Circular dichroism spectra were recorded on a Jasco J-1500 CD Spectrometer equipped with a thermoelectrically controlled cell holder using a quartz cell with a 1.0 mm optical path length. Purified proteins at 0.05 mg/mL were dialyzed into 10 mM Phosphate buffer pH = 7.3 with 150 mM NaCl and scanned between wavelengths from 170 to 260 nm at 25°C. The molar ellipticity from three scans were averaged to provide the final curves presented. For the melt curve analysis, the REPO-NAB protein was continuously scanned over a temperature range from 25°C to 98°C increasing a rate of 0.1°C/min.

### RoseTTAFold and AlphaFold3 structural predictions

YY1’s full amino acid sequence was submitted to RoseTTAFold and AlphaFold3 with default parameters.

### Computational analyses of YY1 genomic binding sites

Data from previous studies (27, 34, 35) and ENCODE (33) were used to generate the Venn diagram depicting YY1’s adherence to the transcription factor binding paradox. HOMER (36) was used to identify the top CLIP-seq motifs for YY1 by using the command findMotifsGenome.pl with the following parameters: -size 100 bp -rna -noknown -nocheck -len 12,18, 25, 30 -mis = 3 -bg BirA_BothStrands_CTRL_CLIP with this file representing background CLIP-seq reads from a BirA pulldown from a cell line with untagged YY1. CompareMotifs.pl was then used to reach a final list of 22 motifs in which the top 6, which represented ∼10% of YY1 CLIP-seq binding sites, were used for further analysis. Two of these sequences are used within the project as “Bioinformatically Derived Sequence 1 and 2” (Figure S3B). Next, we used the JASPAR web database (MA0095.4) to identify genome wide YY1 cognate binding motifs within mm10 and JM8.N4 mESC YY1 CHIP-seq data was obtained (34). Finally, we utilized ENCODE E14.5 ATAC data (33) to identify accessible regions of the mouse genome. We intersected all three previously mentioned datasets with this ATAC dataset to selectively choose motifs and binding sites that are within E14.5 mESC accessible regions. After these initial intersections, we used bedtools -intersect to identify overlapping regions across the datasets and plotted the number of overlapping regions in R using eulerR.

## RESULTS

### The canonical DNA binding domain of YY1 has a higher affinity for ssRNA than dsDNA

To understand the nucleic acid binding properties of YY1 and reconcile the divergent reports of which regions of the protein are responsible for RNA binding activity (26, 27), we designed, expressed, and purified YY1 fragments, then utilized them in quantitative fluorescence polarization assays with a panel of model nucleic acid binding partners. Although denaturing preparations of the protein have been widely used in the past (16, 25, 27, 37) we were concerned that specific activity variation from the refolding process could compromise the strength of conclusions that could be drawn (including incomplete binding saturation in affinity assays). To avoid these potential complications, we chose native purification conditions for full length and fragments of YY1 (Figure S1); only modest activity variation amongst preparations and complete saturation in binding affirmed consistent and high specific activity. Our panel of nucleic acid substrates (Figure 2A) spans the range of strong to weak binding interactions for YY1 reported in the literature (25, 27, 35). For cognate dsDNA reported to have tight binding, we screened the YY1 consensus sequence defined by SELEX (38), as well as the AAVP5 element which was co-crystallized with the sole known DNA binding domain of YY1 (25, 37), its 4 C-terminal C_2_H_2_ zinc finger (ZnF module) (Figure 1A). As a further comparison point, we employed a mutated variant of a 30-bp duplex element from the *Rpl30* promoter, one of the many essential genes regulated by YY1 (18, 39). *Rpl30*’s promoter contains a consensus sequence identical to that defined by SELEX, and it has been previously shown that scrambling the YY1 binding site within this promoter dramatically erodes YY1 affinity (27). As the unmutated *Rpl30* duplex displayed binding affinity slightly tighter than the SELEX consensus sequence for full length YY1 (Figure S2, S3A), we present only the *Rpl30* scrambled mutant data in Figure 2. Both CLIP-seq (27) and CLAP-seq (40) indicate that there is little RNA-sequence motif selectivity in living cells, and prior *in vitro* qualitative studies detected little apparent specificity (26), apart from a bias for U-over C-rich sequences (27, 29). To sample sequence, structure, and a small range of lengths, we assembled a panel of RNA oligonucleotides to probe YY1 RNA-binding affinity. For most YY1 constructs, we relied on the previously described *ARID1A* RNA species as a positive control for RNA binding (27, 41) and as a putative negative control, we chose an RNA devoid of secondary structure and monotonic in sequence, polyuridine (24U), but other RNA oligonucleotides were screened with some protein fragments (Figure S3B, S4).

**Figure 2.**
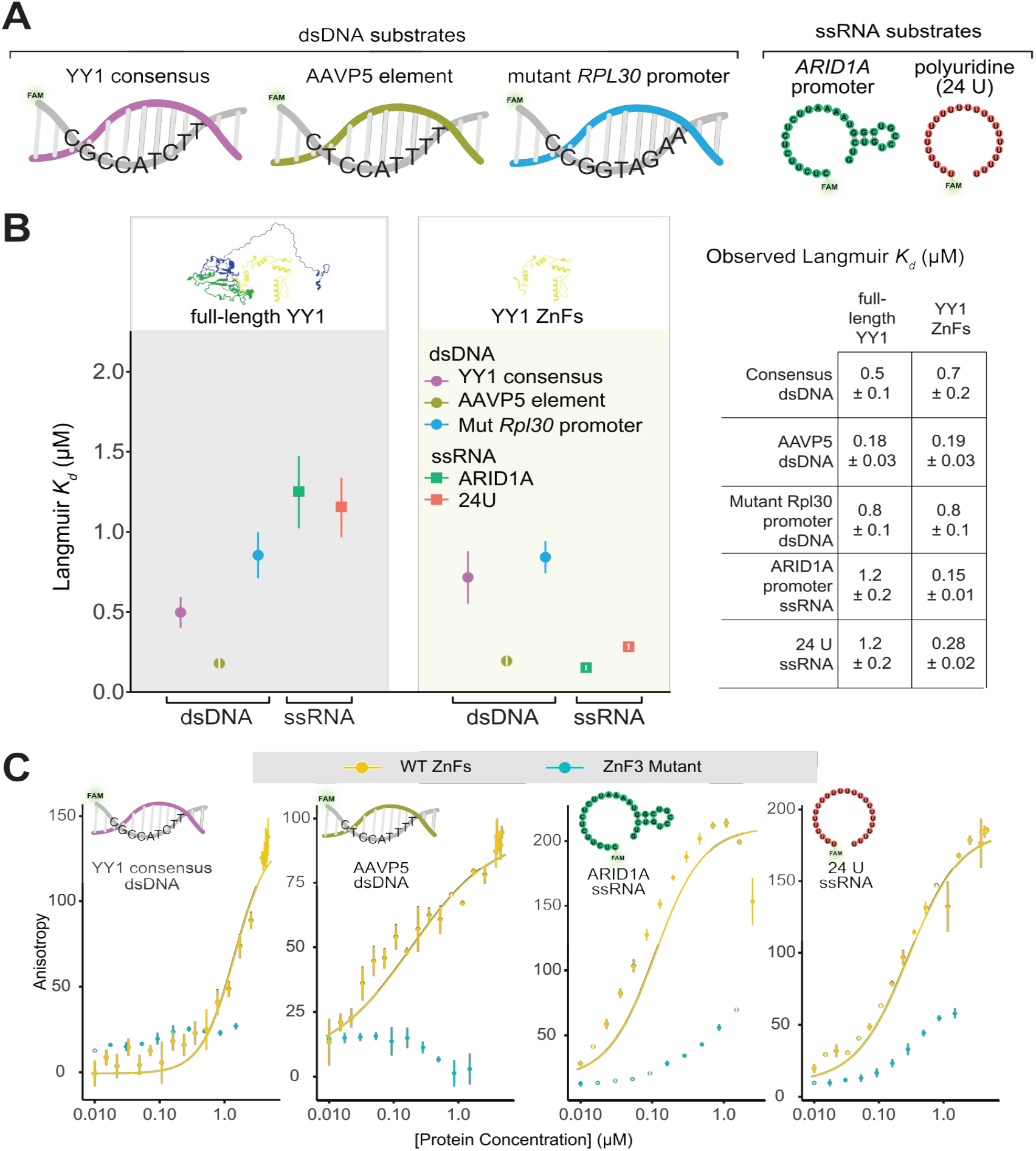
DNA and RNA binding of full length YY1 versus its zinc finger module. **(A)** 5’ FAM labeled nucleic acid panel used in fluorescence polarization assays. ssRNAs are depicted as their predicted RNA structure when input into RNAfold (42) with default parameters. **(B)** Plot of the measured binding affinities (observed Langmuir monovalent binding isotherm *K*_*d*_) of full length YY1 and its 4 C_2_H_2_ zinc fingers (ZnFs, AA 293-414) for our panel of nucleic acids. Error reported represent the standard error within the fit. (Right) Table with numeric values for measured *K*_*d*_ values. **(C)** Fluorescence polarization binding curves comparing WT ZnFs to the ZF3 mutant (AA 365-369 to alanine) for our panel of nucleic acids. Error bars represent the standard deviation from 3 technical replicates, for clarity, only the Hill-equation fits are presented.

Purified full-length YY1 demonstrated expected binding affinities for our panel of nucleic acids (Figure 2A and B, Figure S2), in excellent accordance with previously observed *K*_*d*_ values (27, 37, 43), and overall displayed higher affinity for dsDNA substrates compared to ssRNA substrates. We then purified and assayed the ZnF module of YY1 against our panel of nucleic acids. Although it has been previously reported that the zinc fingers of YY1 can non-specifically bind RNA (26) we were surprised to discover the strength of these interactions exceeded the affinities for dsDNA substrates within our panel by several-fold (Figure 2B). Direct comparisons of ssRNAs with the sequence and length corresponding to the transcribed products of various DNA template strands within our panel, also demonstrated several-fold higher affinity (Figure S4B), excluding sequence- and length-effects as possible explanations for tighter RNA binding. Consistent with previous observations (26, 27), the ZnF module displayed little sequence- or length-specificity (Figure S4A) for RNA, nor any apparent preference for the RNA’s capacity to adopt stable structure (Figure 2A, B, S3B, and S4). We note that prior measurements did not perform direct comparisons of DNA and RNA binding of the ZnF module to the full-length YY1 protein, underscoring the value of our systematic comparisons to reveal potentially important differences.

The capacity of YY1 to bind both DNA and RNA raises the question of whether both nucleic acids bind a similar interface within the ZnF module. Others have noted competitive binding of RNA and dsDNA with the ZnF module (26) and full length YY1 (29, 30), but the details of this interface on the protein side remain obscure. To further probe whether DNA and RNA bind an overlapping interface, we mutated residues S365-N369 to alanine, targeting residues in ZnF3 that participate in both specific base and nonspecific phosphate backbone interactions with the central CATT motif of the consensus sequence (25), while preserving cysteine and histidine residues requisite for C_2_H_2_ zinc finger folding. As anticipated, these mutations ablated apparent dsDNA binding (Figure 2C), yet they also dramatically attenuated ssRNA binding (Figure 2C) supporting the hypothesis that both types of nucleic acids may compete for the same interface within the ZnFs. We note that the complete ablation of dsDNA affinity by this pentamutant is distinct from the severe, but still detectable, erosion of RNA affinity, suggesting overlapping, but non-identical interfaces within the zinc finger module for DNA and RNA binding. Consonant with this interpretation, a previous NMR titration of ssRNA suggested that ZnF1-2 account for most of the RNA-binding chemical shift perturbation (26), and while there is some overlap with sidechains involved in DNA binding in these first two fingers (25), there are seemingly distinct surfaces involved in RNA binding ZnF module.

Intriguingly, while our DNA binding affinity measurements for the ZnF module are similar to those previously observed (27, 37, 43) (AAVP5 dsDNA *K*_*d*_ measurements range from 0.47 ± 0.05 µM (37) to 0.58 ± 0.04 µM (43), as compared to our value of 0.18 ± 0.03 µM), we find the RNA-binding affinity of full-length YY1 protein is an order of magnitude lower than that of its C-terminal ZnF-module. This suggests the remaining portion of the protein may in some way interfere with this domain’s intrinsic RNA-binding properties.

### The conserved REPO domain of YY1 binds nucleic acids when isolated from the rest of the N-terminal domain’s repression

We sought to further biochemically dissect the regions of the protein N-terminal to the ZnFs in order to delineate binding activity modulation properties and/or other potential nucleic acid binding domains. We approached the design of our first N-terminal protein construct by simply dividing YY1 into two fragments: the ZnF module (Figure 2), and the remaining N-terminal portion of the protein (amino acids 1-297) (Figure 3A, N-terminus). Previously Sigova et al. attributed RNA binding to this identical region of YY1 (27), which is consistent with previously noted flexible, nonspecific, electrostatically-driven interactions that IDRs can partake in (44, 45). This hypothesis is further supported by the presence of two RNA-binding arginine rich motifs (ARMs) within the N-terminus; an amino acid motif that has been proposed to be a general feature of TFs that bind RNA to promote their proper genomic localization (28). Yet after purifying and assaying the binding capacity of this N-terminal fragment of YY1, we were surprised to observe that this protein fragment had little apparent affinity for our panel of nucleic acids (Figure 3B).

**Figure 3.**
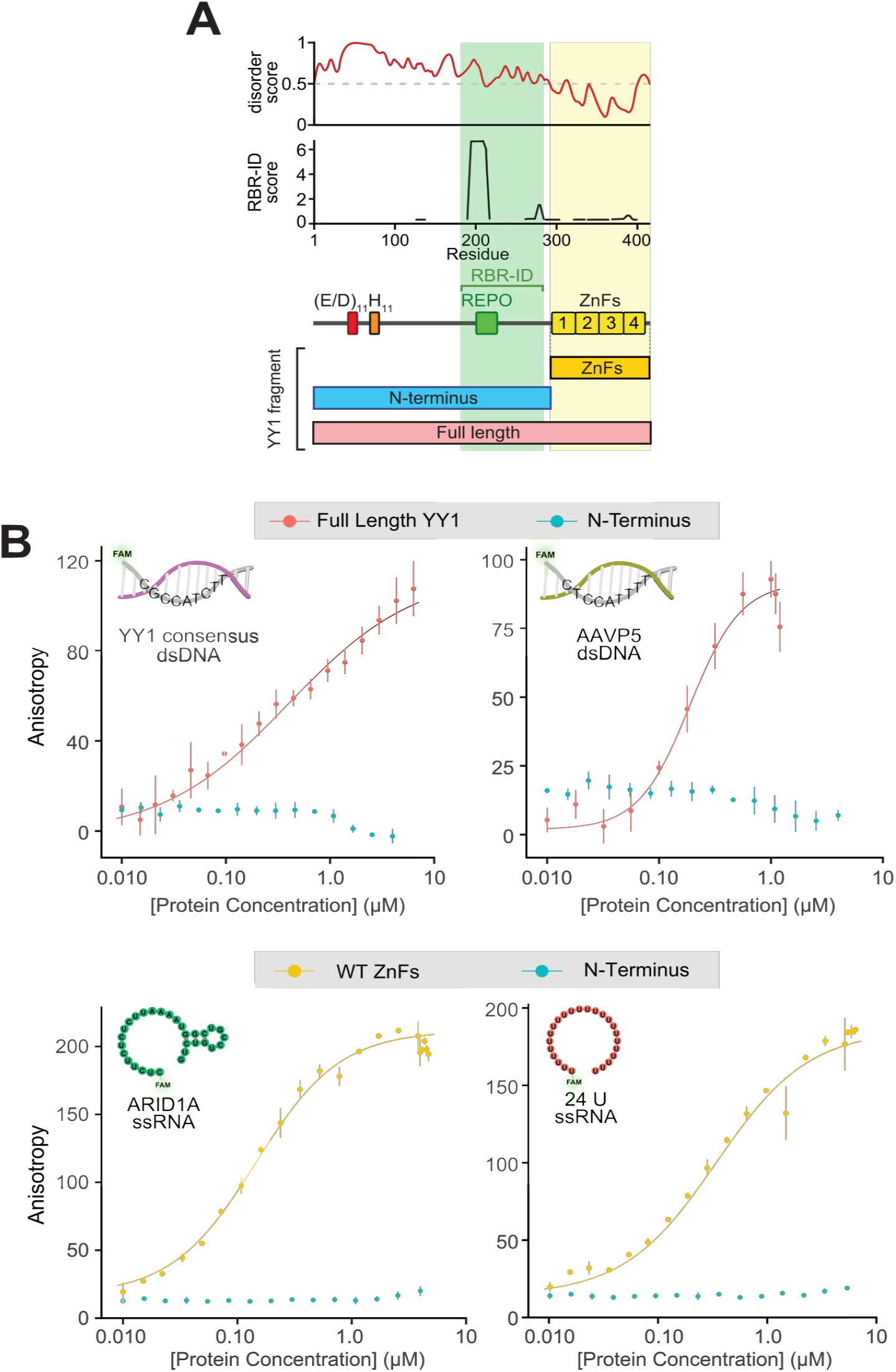
YY1’s N-terminus (AA 1-297) does not bind nucleic acids. **(A)** YY1 domain structure map. (Top) IUPRED3 disorder prediction spanning YY1. (Middle) RNA Binding Region (RBR)-ID score (46) displaying the amino acid residues significantly cross-linked to nuclear RNA within E14 mESCs. (Bottom) Protein constructs that have been purified and assayed in this figure. **(B)** Fluorescence polarization binding experiments for the indicated protein constructs and nucleic acids. Error bars represent standard deviation across three technical replicates.

To reconcile the absence of apparent nucleic acid binding by the N-terminus with the prior study that noted weak binding (27), we sought additional pieces of evidence that could aid the design of further subdivisions of the N-terminus. RBR-ID mass spectrometry data (46) suggests a region of the N-terminus roughly corresponding to the REPO domain of YY1 is significantly crosslinked to nuclear RNA within mouse embryonic stem cells (Figure 4A). The REPO domain has been shown to be both necessary and sufficient for recruitment of Polycomb group proteins to DNA, resulting in transcriptional silencing at loci of recruitment (21) and is highly conserved at the sequence level (82% identity) to the *Drosophila* homolog of YY1, PHO. With this information, we expressed and purified a protein construct with 20 amino acids flanking the two main RBR-ID peaks (Figure 4A, REPO-NAB, AA 164-301) and assessed whether it could bind our panel of nucleic acids. Surprisingly, we observed both DNA and RNA binding capacity, leading us to call this fragment the REPO-NAB for its *<u>n</u>*ucleic *<u>a</u>*cid *<u>b</u>*inding (Figure 4B). REPO-NAB DNA binding was unanticipated, as this region of the protein lacks homology to DNA-binding folds and the only annotated DNA-binding portion of the protein is the C-terminal four C_2_H_2_ ZnF module.

**Figure 4.**
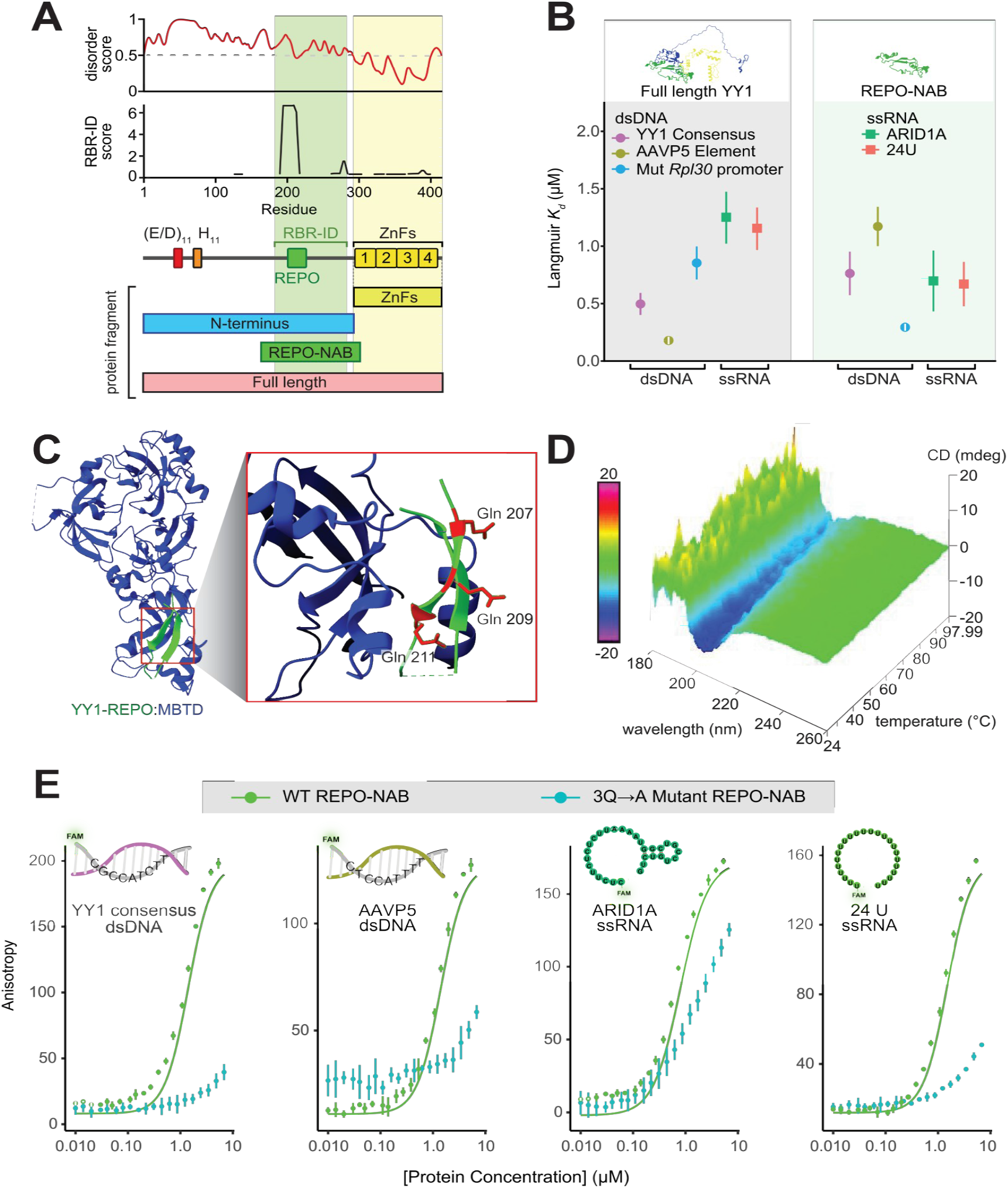
The REPO-NAB domain of YY1 maintains secondary structure in isolation and can bind nucleic acids. **(A)** YY1 domain structure map displaying the spans of YY1 protein constructs used, including the REPO-NAB (AA 164-301; spanning two RBR-ID peaks with 20 amino acid cushion). (Top) IUPRED3 disorder prediction spanning YY1. (Middle) RNA Binding Region (RBR)-ID score (46) displaying the amino acid residues significantly cross-linked to nuclear RNA within E14 mESCs. (Bottom) Protein constructs that have been purified and assayed in this figure as compared to those presented in Figure 3. **(B)** Langmuir *K*_*d*_ plot displaying the measured binding affinities of full length YY1 and the REPO-NAB fragment for our panel of nucleic acids. Error bars represent the standard error of the fit. **(C)** YY1-REPO:MBTD crystal structure (PDB: 4C5I) (50). (Inset) Zoom in of the YY1:MBTD protein-protein interface with Glutamines 207, 209, and 211 depicted in red. **(D)** Temperature dependent circular dichroism of purified REPO-NAB domain indicative of β-sheet character. **(E)** Fluorescence polarization binding curves comparing WT REPO-NAB domain (red) to the 3Q→A Mutant (Q207A, Q209A, and Q211A) REPO-NAB domain (blue) for the indicated nucleic acid substrates. Error bars represent standard deviation across three technical replicates, for clarity, only the Hill-equation fits are presented, Langmuir fits are available in (Figure S5).

The specificity properties of the REPO-NAB are intriguing: although there is little apparent specificity amongst our panel of RNA species, the DNA-binding selectivity is quite distinct from the ZnF module and the full protein. While the REPO-NAB and full length YY1 display similar affinity for the SELEX consensus motif, the AAVP5 sequence is bound >5 fold more weakly by the REPO-NAB (Figure 4B). The REPO-NAB construct displays a several fold-higher affinity for the mutated *Rpl30* dsDNA promoter element (0.30 ± 0.03 µM), whereas 0.8 ± 0.1 µM for both the full-length protein and the ZnFs (Figure 4B). Collectively these data suggest that the full-length protein suppresses these apparent DNA binding preferences of the REPO-NAB in addition to RNA-binding inhibition.

Although the REPO-NAB is predicted to be intrinsically disordered by IUPRED (47) (Figure 4A), the apparent selectivity of its DNA-binding suggest some degree of structure. We noted that when generating *de novo* structural models of YY1, RoseTTAFold (48) and AlphaFold3 (49) consistently predicts that the REPO domain of YY1 maintains its anti-parallel β-sheet character within the overall structure of the protein. This is consistent with (and likely informed by) a previous crystal structure of the human YY1 REPO domain in complex with the 4MBT domain of human MBTD1, homologous to the Drosophila PhoRC-polycomb repressive complex (50) (Figure 4C). However, it is unknown whether the REPO domain maintains this anti-parallel β-sheet character in isolation. To address this, we performed circular dichroism with biochemically purified REPO-NAB and observed that the anti-parallel β-sheet character of this domain is maintained in isolation although the melting transition is not sharply defined (Figure 4D). Similar β-sheet character, previously noted for the full N-terminus (51), is at least in part attributable to the REBO-NAB. We noted that three glutamines (Q207, Q209, and Q211) were not involved in the YY1:MBTD1 interface and therefore could play a role in the REPO domain’s nucleic acid binding (Figure 4C), given the propensity for glutamines to engage in nucleic acid-recognition (52–55). Mutating each of these three residues to alanine, we achieved a partial separation-of-function mutant without impacting the β-character (Figure S5B): dsDNA binding was ablated, whereas RNA binding attenuated within our focused panel of nucleic acids (Figure 4E).

### Defining the autoinhibitory function of YY1’s N-terminus

A possible explanation for the distinct nucleic acid binding preferences of the REPO-NAB and ZnF module, relative to the full-length protein, is that the N-terminus of YY1 could regulate them. From our binding assays with the N-terminus (AA 1-297, Figure 3) and REPO-NAB (AA 164-301, Figure 4), we can conclude that when the REPO-NAB is covalently linked to the rest of the N-terminus it becomes inhibited from binding any nucleic acid species within our panel. Similarly, the ZnF module’s RNA binding capacity is reduced in the context of the full protein (Figure 2B). To test this hypothesis directly, we performed fluorescence polarization competition experiments assaying the affinity of the ZnFs for the *ARID1A* RNA and the YY1 consensus dsDNA sequence in both the presence and absence of purified N-terminus (Figure 5A and B). For both nucleic acid types we observed decreased binding with the addition of the N-terminus as a function of its concentration. As the N-terminus does not display any detectable nucleic acid binding in the concentration regime (Figure 3B), these observations lead us to believe that the N-terminus can inhibit nucleic acid binding of the ZnFs in *trans i*.*e*., not covalently bound to the ZnFs. While Figure 5A is consistent with the apparent *cis*-attenuation of RNA-binding of this protein region in the context of the full-length protein (compared to ZnF module alone), the *trans* inhibition of DNA binding by the N-terminus is unexpected (Figure 5B), as these properties did not markedly differ between full length YY1 and the ZnF fragment.

**Figure 5.**
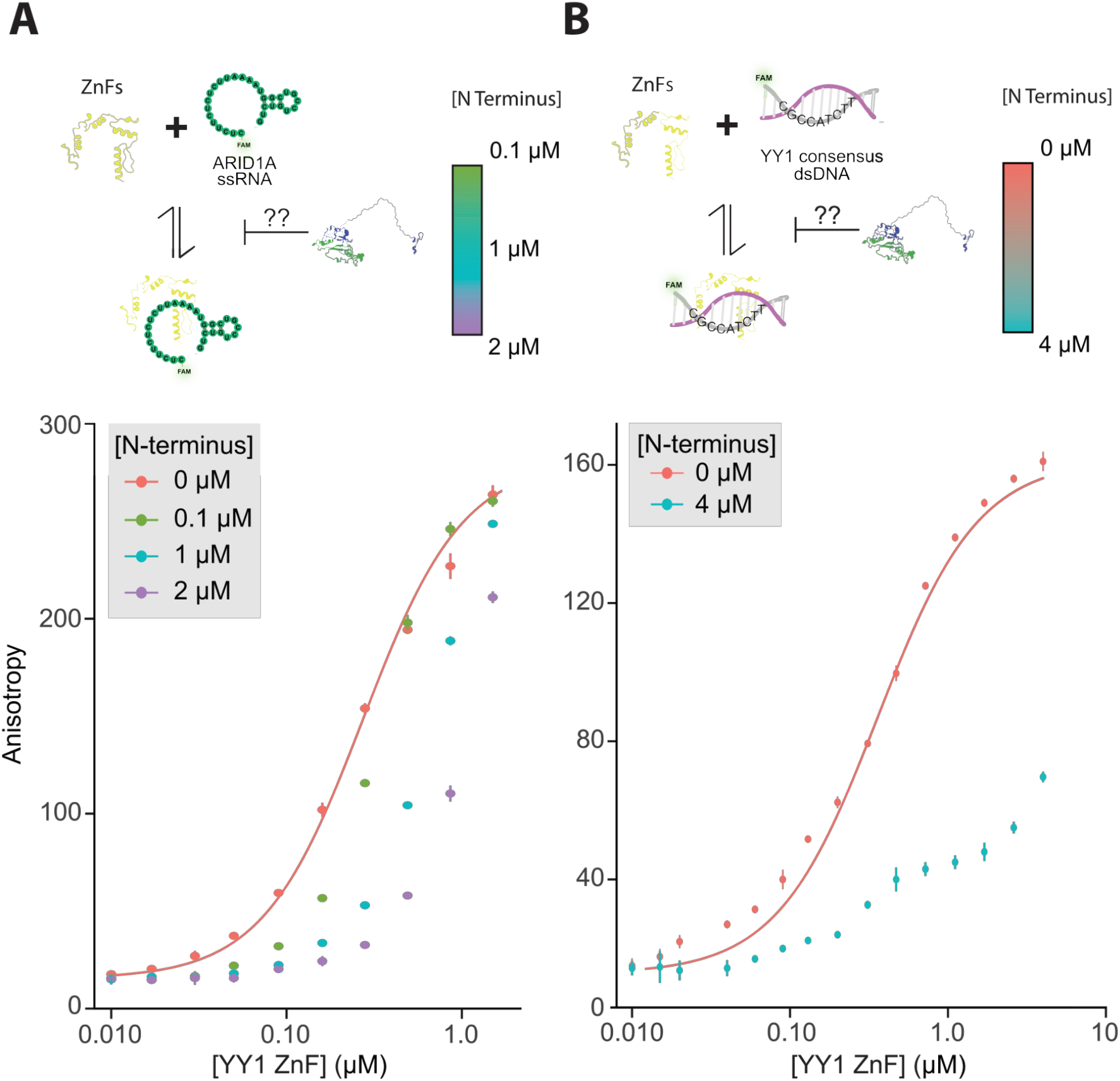
YY1’s N-terminus modulates nucleic acid binding of its ZnF module. **(A** and **B)** Fluorescence polarization competition experiments. The indicated amounts of purified N-terminus were added in *trans* to a ZnFs titration of each of the representative nucleic acid substrates indicated. Error bars represent standard deviation across three technical replicates.

We hypothesized that intramolecular inhibition could occur via charge complementarity between nucleic acid binding domains and a stretch of negatively charged amino acids (E/D) within the N-terminal domain of YY1. A portion of YY1’s “core IDR” (cIDR) contains a consecutive stretch of 11 acidic amino acids (AA E43-D53) that are fully conserved across metazoans and have recently been shown to be a component of the cIDR which is both necessary and sufficient to drive phase separation of YY1 in both *in vitro* and cell-based assays (56). This stretch of negative amino acids has also been hypothesized to aid in YY1’s genomic target search by autoinhibition of spurious nucleic acid binding (57). Upon generating an N-terminus construct with these 11 residues (E43-D53) mutated to alanine (Figure S6A), we observed that this mutant, like the wild type N-terminus could not bind our panel of nucleic acids (Figure S6B). Although the REPO-NAB domain is not sensitive to regulation by the E/D patch in *cis*, we sought to perform similar *trans* complementation experiments with the ZnF module and the N-terminus bearing the E/D patch mutation. In this context as well, there was also little difference between the N-terminus and the mutant—both similarly impaired RNA binding of the zinc fingers in *trans* (Figure S6C). Collectively these experiments suggest that rather than localized to a single cluster of the 11 E/D residues that we targeted, the nature of inhibition may be more complex. Charge complementarity-based regulation, if it indeed exists, must be dispersed across the IDR. This E/D cluster accounts for only 11 of the 38 acidic residues in this region of the protein and similar delocalized properties have been noted in other cases (5, 6, 9).

To further investigate this inhibition, we sought to purify a truncated N-terminus lacking the REPO-NAB domain (Figure S7A, AA 1-163) and assess its inhibitory properties. This fragment displayed several orders of magnitude weaker binding to nucleic acids (Figure S7B)— although it was not as binding deficient as the full N-terminus construct, we were unable to measure dissociation constants for any of the nucleic acid species within our panel. Since we observed close to no change in anisotropy for the disordered N-terminus in the concentration range in which the ZnF module reaches saturation with our panel of nucleic acids (0 – 2 µM), we decided to stage competition experiments in order to assess whether the disordered N-terminus could regulate nucleic acid binding. We observed little to no inhibition of binding of either nucleic acid type when we added the disordered N-terminus *in trans* to the ZnFs at concentrations below (Figure S7 C,D), but approaching regimes where this module itself could directly bind (Figure S7B). Considering the discrepancy between the affinities the ZnF module and full length YY1 exhibit for RNA binding, these observations suggests that direct linkage of the REPO-NAB is necessary to orient the N-terminus to inhibit RNA-binding of the ZnF module.

## DISCUSSION

In this work, we systematically interrogate the nucleic acid-binding domain structure, and intrinsic regulation thereof, by other portions of YY1, revealing a surprisingly complex architecture of autoregulatory logic in the absence of additional co-factors. First, we address the controversy regarding which portions of the YY1 protein are responsible for RNA-binding via systematic side-by-side equilibrium binding studies, whose quantitative nature proved essential to our proposed resolution. Semi-quantitative measurements have been used to suggest that the N-terminus of the protein, but not the C-terminal ZnF module, bears weak RNA-binding capacity relative to the full-length protein (27). Whereas more quantitative examinations of the C-terminal ZnFs in isolation, absent data presented for the full protein (26) or the N-terminal portion (26), argue that this module harbours RNA binding capacity thought to be important for function, without clear exclusion of the N-terminus contributions. Consonant with the latter two papers, we find that indeed the C-terminal ZnF module has both DNA- and RNA-binding capacity, but to our surprise, the RNA binding affinity of the ZnF module in isolation exhibits an order of magnitude tighter binding than what we observed with the full-length protein, under identical conditions, whereas the dsDNA affinities are quite similar for both (Figure 2B). We detect no RNA affinity for an identical N-terminal fragment of the protein at a concentration regime more than an order of magnitude higher than where Sigova and colleagues detected weak N-terminal binding (27) (Figure 3B). Remarkably, dissection of the N-terminal domain into the N-terminal IDR (AA 1-163) and the REPO-NAB (AA 164-301) reveals the latter’s marked affinity for RNA (as well as DNA) (Figure 4B) is seemingly suppressed by the former (Figure 3B). It is possible that in the prior work (27), proteolytic fragments bearing the REPO-NAB separate from the N-terminal IDR are present in the preparation of the N-terminal fragment which could account for apparent binding by EMSA and is consistent with the complex pattern of shifted bands observed. Our data suggests both seemingly divergent prior RNA-binding attributions to be true in the sense that there are regions in both fragments that have RNA-binding capacity, and in so doing, we reveal a potent RNA-binding inhibitory property of the N-terminal IDR. This inhibitory property of the N-terminus is not limited to the REPO-NAB--we have discovered a similar regulatory mechanism the N-terminus imposes upon the ZnF module as well. Although our study is limited to the scope of nucleic acid species surveyed and by the *in vitro* nature of our quantitative assays, these observations of the protein’s intrinsic properties may provide insight into the highly context-dependent nature of YY1’s activities *in vivo*.

### The interfaces of RNA and DNA binding overlap in both nucleic acid binding domains

With recent work shedding light on the capacity of TFs to bind both duplex DNA and RNA (28, 58–60), inquiry into the mechanisms and functional consequences of these properties are of great interest. We and others (18, 26–29), have investigated whether dsDNA and ssRNA compete for a similar binding interface within full length YY1. In the most mechanistically detailed prior study, Wai and colleagues (26) used ^15^N-heteronuclear single-quantum-coherence (^15^N-HSQC) NMR experiments to identify amino acid residues in the ZnF module that engaged in RNA binding, finding the bulk of RNA interactions localized to ZnF1 and ZnF2 (26). Although a number of residues that displayed significant chemical shift perturbations in the presence of RNA are involved in dsDNA contacts in the YY1 ZnF module crystal structure (25), the authors noted a contiguous surface of amino acids within ZnF2 (V324, H325, V326, L340, Q344) that was completely distinct from the dsDNA binding interface (26). Mutagenesis of these amino acids to alanine diminished apparent binding to RNA by at least an order of magnitude (26), consistent with this unique interface representing an energetically consequential portion of the RNA-binding capacity. However, retention of DNA-binding affinity of this pentamutant was not evaluated, so it remains uncertain whether it represents a *bona fide* separation of function mutant.

In an attempt to design the converse separation of function mutation, wherein DNA-binding affinity of the ZnF module was selectively perturbed with minimal RNA-impact, we targeted amino acids in a different segment of the ZnF module (AA 365-369, ZnF3, Figure 2C). This mutant indeed displays no measurable DNA binding activity and its RNA-binding was severely attenuated, but still detectable (Figure 2C). This impact on RNA-binding was unexpected, as these residues, and ZnF3, overall exhibit minimal chemical shift perturbations in the presence of RNA (26). One possible explanation for this apparent disparity in the importance of the ZnF3 for RNA binding is the identity of the RNA in our experiments is distinct—the *K*_*d*_ measurements for ssRNA in this prior work range from 20-60 fold weaker than our measurements. The Wai paper used a 14-mer of RNA isolated from a SELEX experiment for their affinity and structural studies (*K*_d_ = 3.8 ± 0.6 µM, although this was within error of a poly-A substrate of the same length in their experiments). Whereas the majority of our experiments were conducted with longer RNA species, we note serial truncation of the ARID1A 30-mer down to a 14-mer element leads to a ∼3-fold affinity loss monotonically across the series (Figure S4A), and this *K*_*d*_ = 0.29 ± 0.1 µM is comparable to two 12-mers of completely different sequence composition (Figure S3B). Thus, size and sequence do not account for our order of magnitude higher affinity measurements for the ZnF module binding RNA in this regime. The methods of protein preparation are highly analogous, the binding buffers are only subtly different, and although the measurement methods differ, both microscale thermophoresis and fluorescence polarization are solution-based measurements free of confounding surface effects. It may be that for shorter ssRNA, the interface of the YY1 ZnF module species less extensively engages ZnF3 and ZnF4. This is consistent with the origin of the sequence used in NMR structural studies, which was selected for using a Zn-coordinating mutant in ZnF4 that should unfold it (26). Nevertheless, our data with longer RNA species and the ZnF3 mutant clearly indicates that this region of the protein can also be important for RNA binding.

We have identified a similar scenario in which DNA and RNA share an overlapping binding interface in the elucidation of the REPO-NAB’s nucleic acid binding properties. Our mutant REPO-NAB, harbouring alanine substitutions at Q207, Q209, and Q211, loses its capacity to bind dsDNA substrates (Figure 4E) but these mutations attenuate, yet do not completely diminish, the ability of this mutant to bind our ssRNA substrates. Recent work implicates ARM-motifs, basic residue rich patches directly adjacent to canonical DNA binding domains of several transcription factors, in promoting their chromatin association through RNA binding (28), a mechanism first proposed for YY1 (27). Although the REPO-NAB does contain a basic patch directly flanking the ZnF module (AA 281-288, Figure S6A), this region is not able to bind nucleic acid in the context of the full N-terminus. While it is possible that this motif does participate in RNA binding as proposed for other ZnF TFs (28), our mutagenesis of the distinct REPO domain within this construct (AA 201-226) suggests that the ARM motif is not playing a dominant role in the nucleic acid binding we observe. Further *in vivo* experiments elucidating how the binding of one nucleic acid type (in this case ssRNA) and the loss of binding for another (dsDNA in this case) can affect the context-dependent localization and function of full length YY1 can be undertaken with separation-of-function mutants of the sort we report here. The identification of the REPO-NAB’s nucleic acid binding properties also affords another example to the growing list of cryptic/ancillary nucleic acid binding domains (10, 58, 61). Understanding what this additional binding capacity imparts upon their respective proteins in cellular contexts is the next key step to define their functional importance.

### YY1’s N-terminus tunes the nucleic acid binding affinity of the REPONAB and the ZnF module

Here we begin to dissect the autoinhibitory properties of the N-terminus of YY1, in both intra- and inter-molecular mechanisms (*cis* versus *trans*) on the two nucleic acid binding domains of YY1. Our measurements reveal how the N-terminus, either covalently attached in single polypeptide or added in *trans*, affects nucleic acid binding of the REPO-NAB and ZnF modules. We find that when the REPO-NAB is physically linked to the remainder of the N-terminus of YY1 (the N-terminal IDR), it is efficiently inhibited from binding both ssRNA and dsDNA (Figure 3). Similarly, in competition experiments assaying the ZnF module’s capacity to engage nucleic acids in the presence/absence of the N-terminus we observe a drastic inhibition of either type of nucleic acid binding. The impact on DNA binding is unexpected as we envisioned that reconstituting the inhibitory interaction in *trans* would begin to recapitulate the properties of full length YY1—blunting of the intrinsic capacity of the ZnF module’s capacity to bind RNA efficiently, while preserving the DNA-binding capacity. However, adding these two fragments in *trans* may fail to capture some aspect of orientation provided by direct linkage in the native full protein. The differences of the N-terminus added in *trans* versus *cis* could be interpreted as revealing the potential for regulation by YY1’s reported capacity to dimerize/multimerize (17, 18, 20, 62) in a potentially RNA-dependent manner (17, 18, 62). One possible function of these mechanisms is to keep YY1 in an inhibited state, awaiting further protein/nucleic acid interactions to ultimately impart context-dependent functionality. Further experiments will be needed to precisely map the interfaces between the REPO-NAB, the ZnF module and the N-terminus that impart this nucleic acid binding inhibition within YY1, and to determine how generalizable these regulatory features are to other transcription factors, that often have IDRs (5, 6, 8, 9) and poorly understood additional RNA-binding regions (28).

### How the two newly identified YY1 activities may relate to its myriad functions

Given the wide spectrum of functions ascribed to YY1 and the large collection of well-established protein binding partners in each of these scenarios (18–23), we propose that the intrinsic YY1 nucleic acid binding and autoregulation thereof could be key components of these diverse regulatory outcomes. The acidic transactivation domain of YY1 has been mapped in cell-based reporter assays to the N-terminus, with the precise boundaries of this region differing somewhat but generally including amino acids 1-99 (15). This constitutes the bulk of the N-terminal IDR construct (Figure S6A) which inhibits nucleic acid binding of the REPO-NAB and is necessary but not sufficient for ZnF module RNA-binding suppression in *cis*. Thus, the act of participating in transcriptional activation interactions (4), could relieve nucleic-acid binding inhibition of the REPO-NAB and enable RNA-binding of the ZnFs. Similarly, a core region of the N-terminal IDR spanning amino acids 43-80 has recently been noted to drive phase separation *in vitro* with corresponding impact on cellular YY1 function (56). These properties require 11 consecutive histidine residues within this core IDR which have also been proposed to mediate the homodimerization/multimerization of YY1 via zinc coordination (63). Interestingly, RNA has been proposed to facilitate the YY1 dimerization (18, 62) thought to be essential for through space architectural linkage of two YY1 DNA binding sites (18). Given our data that suggests this YY1 region is involved in suppressing RNA-binding of either of the two nucleic acid binding domains, we postulate that the YY1 protein-protein interface can either act to dimerize the protein or inhibit RNA-binding in *cis*, and that the addition of RNA competitively releases this N-terminal inhibitory region to engage in chromosome looping *trans*-YY1 interactions. This is consistent with a prior observation that Rbm25 stabilizes YY1 binding to chromatin via protein-protein interactions, and some of this stabilization could be accounted for by YY1’s RNA binding (64).

More broadly, the putative interplay between protein-partner binding and RNA/DNA binding could account for the remarkable diversity of YY1 cellular functions. Specific and nonspecific nucleic acid binding by the domains that we have interrogated could play a role in YY1’s genomic localization, supporting the observations of pervasive transcription factor “trapping” posited by others (27, 28) and implicating IDRs in proper TF binding site engagement (5, 8, 9, 56, 57). Our work has demonstrated that rigorous, systematic, biophysical approaches can uncover unannotated properties of very well-characterized proteins and can therefore guide further investigation to their function. We contend that defining the intrinsic properties of the YY1 polypeptide with respect to nucleic acid binding and its autoinhibition, represents a critical advance to elucidating YY1’s precise molecular mechanisms and their functional impact in transcription, repression and genome architecture. In future studies, we hope to characterize the interplay of these features with the catalogue of known YY1 protein binding partners, as well as determine the functional consequences of these newly defined properties *in vivo*.

## DATA AVAILABILITY

The data underlying this article are available in Zenodo at 10.5281/zenodo.13694509.

## SUPPLEMENTARY DATA

Supplementary Data are available at NAR online.

## AUTHOR CONTRIBUTIONS

Jimmy Elias: Conceptualization, Methodology, Investigation, Formal analysis, Validation, Supervision, Writing—original draft. Alex Ruthenburg: Funding Acquisition, Project Administration, Conceptualization, Visualization, Writing—review & editing. Jane Rosin: Investigation. Amanda Keplinger: Investigation, Writing—editing.

## ACKNOWLEDGEMENTS

This work was supported by the National Institutes of Health grants R35-GM145373 to A.J.R, 5T32GM007197 (PI: Lucia Rothman-Denes) and 5R25GM109439 (PI: Nancy Schwartz); and a GRFP from the National Science Foundation to A.J.K.

Funding for open access charge: National Institutes of Health.

## REFERENCES

1. Hnisz, D., Abraham, B.J., Lee, T.I., Lau, A., Saint-André, V., Sigova, A.A., Hoke, H.A. and Young, R.A. (2013) Super-enhancers in the control of cell identity and disease. Cell, 155, 934.

2. Takahashi, K. and Yamanaka, S. (2006) Induction of Pluripotent Stem Cells from Mouse Embryonic and Adult Fibroblast Cultures by Defined Factors. Cell, 126, 663–676.

3. Lambert, S.A., Jolma, A., Campitelli, L.F., Das, P.K., Yin, Y., Albu, M., Chen, X., Taipale, J., Hughes, T.R. and Weirauch, M.T. (2018) The Human Transcription Factors. Cell, 172, 650–665.

4. Sanborn, A.L., Yeh, B.T., Feigerle, J.T., Hao, C.V., Townshend, R.J.L., Aiden, E.L., Dror, R.O. and Kornberg, R.D. (2021) Simple biochemical features underlie transcriptional activation domain diversity and dynamic, fuzzy binding to mediator. eLife, 10, 1–42.

5. Jana, T., Brodsky, S. and Barkai, N. (2021) Speed–Specificity Trade-Offs in the Transcription Factors Search for Their Genomic Binding Sites. Trends Genet., 10.1016/j.tig.2020.12.001.

6. Brodsky, S., Jana, T., Mittelman, K., Chapal, M., Kumar, D.K., Carmi, M., Brodsky, S., Jana, T., Mittelman, K., Chapal, M., et al. (2020) Intrinsically Disordered Regions Direct Transcription Factor In Vivo Binding Specificity Article Intrinsically Disordered Regions Direct Transcription Factor In Vivo Binding Specificity. Mol. Cell, 10.1016/j.molcel.2020.05.032.

7. Mazzocca, M., Loffreda, A., Colombo, E., Fillot, T., Gnani, D., Falletta, P., Monteleone, E., Capozi, S., Bertrand, E., Legube, G., et al. (2023) Chromatin organization drives the search mechanism of nuclear factors. Nat. Commun., 14, 6433.

8. Chen, Y., Cattoglio, C., Dailey, G.M., Zhu, Q., Tjian, R. and Darzacq, X. (2022) Mechanisms governing target search and binding dynamics of hypoxia-inducible factors. eLife, 11, e75064.

9. Kumar, D.K., Jonas, F., Jana, T., Brodsky, S., Carmi, M. and Barkai, N. (2023) Complementary strategies for directing in vivo transcription factor binding through DNA binding domains and intrinsically disordered regions. Mol. Cell, 83, 1462-1473.e5.

10. Hansen, A.S., Amitai, A., Cattoglio, C., Tjian, R. and Darzacq, X. (2020) Guided nuclear exploration increases CTCF target search efficiency. Nat. Chem. Biol., 16, 257–266.

11. Panne, D., Maniatis, T. and Harrison, S.C. (2007) An atomic model of the interferon-beta enhanceosome. Cell, 129, 1111–1123.

12. Göös, H., Kinnunen, M., Salokas, K., Tan, Z., Liu, X., Yadav, L., Zhang, Q., Wei, G.-H. and Varjosalo, M. (2022) Human transcription factor protein interaction networks. Nat. Commun., 13, 766.

13. Tycko, J., DelRosso, N., Hess, G.T., Aradhana Banerjee, A., Mukund, A., Van, M.V., Ego, B.K., Yao, D., Spees, K., et al. (2020) High-Throughput Discovery and Characterization of Human Transcriptional Effectors. Cell, 183, 2020-2035.e16.

14. Soto, L.F., Li, Z., Santoso, C.S., Berenson, A., Ho, I., Shen, V.X., Yuan, S. and Fuxman Bass, J.I. (2022) Compendium of human transcription factor effector domains. Mol. Cell, 82, 514–526.

15. Lee, J.-S., See, R.H., Galvin, K.M., Wang, J. and Shi, Y. (1995) Functional interactions between YY1 and adenovirus E1A. Nucleic Acids Res., 23, 925–931.

16. Shi, Y., Seto, E., Chang, L.S. and Shenk, T. (1991) Transcriptional repression by YY1, a human GLI-Krüppel-related protein, and relief of repression by adenovirus E1A protein. Cell, 67, 377–388.

17. Shi, Y., Lee, J.S. and Galvin, K.M. (1997) Everything you have ever wanted to know about Yin Yang 1… Biochim. Biophys. Acta - Rev. Cancer, 1332.

18. Weintraub, A.S., Li, C.H., Zamudio, A.V., Sigova, A.A., Hannett, N.M., Day, D.S., Abraham, B.J., Cohen, M.A., Nabet, B., Buckley, D.L., et al. (2017) YY1 Is a Structural Regulator of Enhancer-Promoter Loops. Cell, 171, 1573-1588.e28.

19. Seto, E., Shi, Y. and Shenk, T. (1991) YY1 is an initiator sequence-binding protein that directs and activates transcription in vitro. Nature, 354, 241–245.

20. López-Perrote, A., Alatwi, H.E., Torreira, E., Ismail, A., Ayora, S., Downs, J.A. and Llorca, O. (2014) Structure of Yin Yang 1 oligomers that cooperate with RuvBL1-RuvBL2 ATPases. J. Biol. Chem., 289, 22614–22629.

21. Wilkinson, F.H., Park, K. and Atchison, M.L. (2006) Polycomb recruitment to DNA in vivo by the YY1 REPO domain. Proc. Natl. Acad. Sci. U. S. A., 103, 19296–19301.

22. Wang, J., Wu, X., Wei, C., Huang, X., Ma, Q., Huang, X., Faiola, F., Guallar, D., Fidalgo, M., Huang, T., et al. (2018) YY1 Positively Regulates Transcription by Targeting Promoters and Super-Enhancers through the BAF Complex in Embryonic Stem Cells. Stem Cell Rep., 10, 1324–1339.

23. Wu, S., Shi, Y., Mulligan, P., Gay, F., Landry, J., Liu, H., Lu, J., Qi, H.H., Wang, W., Nickoloff, J.A., et al. (2007) A YY1-INO80 complex regulates genomic stability through homologous recombination-based repair. Nat. Struct. Mol. Biol., 14, 1165–1172.

24. Zhang, X., Blumenthal, R.M. and Cheng, X. (2024) Updated understanding of the protein–DNA recognition code used by C2H2 zinc finger proteins. Curr. Opin. Struct. Biol., 87, 102836.

25. Houbaviy, H.B., Usheva, A., Shenk, T. and Burley, S.K. (2002) Cocrystal structure of YY1 bound to the adeno-associated virus P5 initiator. Proc. Natl. Acad. Sci., 93, 13577–13582.

26. Wai, D.C.C., Shihab, M., Low, J.K.K. and Mackay, J.P. (2016) The zinc fingers of YY1 bind single-stranded RNA with low sequence specificity. Nucleic Acids Res., 44, 9153–9165.

27. Sigova, A.A., Abraham, B.J., Ji, X., Mollinie, B., Hannett, N.M., Guo, Y.E., Jangi, M., Giallourakis, C.C., Sharp, P.A. and Young, R.A. (2015) Transcription factor trapping by RNA in gene regulatory elements. Science, 350, 978–982.

28. Oksuz, O., Henninger, J.E., Warneford-Thomson, R., Zheng, M.M., Erb, H., Vancura, A., Overholt, K.J., Hawken, S.W., Banani, S.F., Lauman, R., et al. (2023) Transcription factors interact with RNA to regulate genes. Mol. Cell, 83, 2449-2463.e13.

29. Belak, Z.R. and Ovsenek, N. (2007) Assembly of the Yin Yang 1 Transcription Factor into Messenger Ribonucleoprotein Particles Requires Direct RNA Binding Activity*. J. Biol. Chem., 282, 37913–37920.

30. Chen, Z.S., Ou, M., Taylor, S., Dafinca, R., Peng, S.I., Talbot, K. and Chan, H.Y.E. (2023) Mutant GGGGCC RNA prevents YY1 from binding to Fuzzy promoter which stimulates Wnt/β-catenin pathway in C9ALS/FTD. Nat. Commun., 14, 8420.

31. Jeon, Y. and Lee, J.T. (2011) YY1 Tethers Xist RNA to the inactive X nucleation center. Cell, 146, 119–133.

32. Nabeel-Shah, S., Pu, S., Burns, J.D., Braunschweig, U., Ahmed, N., Burke, G.L., Lee, H., Radovani, E., Zhong, G., Tang, H., et al. (2024) C2H2-zinc-finger transcription factors bind RNA and function in diverse post-transcriptional regulatory processes. Mol. Cell, 0.

33. Gorkin, D.U., Barozzi, I., Zhao, Y., Zhang, Y., Huang, H., Lee, A.Y., Li, B., Chiou, J., Wildberg, A., Ding, B., et al. (2020) An atlas of dynamic chromatin landscapes in mouse fetal development. Nature, 583, 744–751.

34. Hsieh, T.-H.S., Cattoglio, C., Slobodyanyuk, E., Hansen, A.S., Darzacq, X. and Tjian, R. (2022) Enhancer–promoter interactions and transcription are largely maintained upon acute loss of CTCF, cohesin, WAPL or YY1. Nat. Genet., 54, 1919–1932.

35. Chen, K., Lu, Y., Shi, K., Stovall, D.B., Li, D. and Sui, G. (2019) Functional analysis of YY1 zinc fingers through cysteine mutagenesis. FEBS Lett., 593, 1392–1402.

36. Heinz, S., Benner, C., Spann, N., Bertolino, E., Lin, Y.C., Laslo, P., Cheng, J.X., Murre, C., Singh, H. and Glass, C.K. (2010) Simple combinations of lineage-determining transcription factors prime cis-regulatory elements required for macrophage and B cell identities. Mol. Cell, 38, 576–589.

37. Houbaviy, H.B. and Burley, S.K. (2001) Thermodynamic analysis of the interaction between YY1 and the AAV P5 promoter initiator element. Chem. Biol., 8, 179–187.

38. Yant, S.R., Zhu, W., Millinoff, D., Slightom, J.L., Goodman, M. and Gumucio, D.L. (1995) High affinity YY1 binding motifs: identification of two core types (ACAT and CCAT) and distribution of potential binding sites within the human β globin cluster. Nucleic Acids Res., 23, 4353–4362.

39. Shrivastava, A. and Calame, K. (1994) An analysis of genes regulated by the multi-functional transcriptional regulator Yin Yang-1. Nucleic Acids Res., 22, 5151–5155.

40. Guo, J.K., Blanco, M.R., Walkup, W.G., Bonesteele, G., Urbinati, C.R., Banerjee, A.K., Chow, A., Ettlin, O., Strehle, M., Peyda, P., et al. (2024) Denaturing purifications demonstrate that PRC2 and other widely reported chromatin proteins do not appear to bind directly to RNA in vivo. Mol. Cell, 0.

41. Heller, D., Krestel, R., Ohler, U., Vingron, M. and Marsico, A. (2017) ssHMM: extracting intuitive sequence-structure motifs from high-throughput RNA-binding protein data. Nucleic Acids Res., 45, 11004–11018.

42. Lorenz, R., Bernhart, S.H., Höner zu Siederdissen, C., Tafer, H., Flamm, C., Stadler, P.F. and Hofacker, I.L. (2011) ViennaRNA Package 2.0. Algorithms Mol. Biol., 6, 26.

43. Golebiowski, F.M., Górecki, A., Bonarek, P., Rapala-Kozik, M., Kozik, A. and Dziedzicka-Wasylewska, M. (2012) An investigation of the affinities, specificity and kinetics involved in the interaction between the Yin Yang 1 transcription factor and DNA. FEBS J., 279, 3147–3158.

44. Shrinivas, K., Sabari, B.R., Coffey, E.L., Klein, I.A., Boija, A., Zamudio, A.V., Schuijers, J., Hannett, N.M., Sharp, P.A., Young, R.A., et al. (2019) Enhancer Features that Drive Formation of Transcriptional Condensates. Mol. Cell, 75, 549-561.e7.

45. Habchi, J., Tompa, P., Longhi, S. and Uversky, V.N. (2014) Introducing protein intrinsic disorder. Chem. Rev., 114, 6561–6588.

46. He, C., Sidoli, S., Warneford-Thomson, R., Tatomer, D.C., Wilusz, J.E., Garcia, B.A. and Bonasio, R. (2016) High-Resolution Mapping of RNA-Binding Regions in the Nuclear Proteome of Embryonic Stem Cells. Mol. Cell, 64, 416–430.

47. Erdős, G., Pajkos, M. and Dosztányi, Z. (2021) IUPred3: prediction of protein disorder enhanced with unambiguous experimental annotation and visualization of evolutionary conservation. Nucleic Acids Res., 49, W297–W303.

48. Baek, M., DiMaio, F., Anishchenko, I., Dauparas, J., Ovchinnikov, S., Lee, G.R., Wang, J., Cong, Q., Kinch, L.N., Schaeffer, R.D., et al. (2021) Accurate prediction of protein structures and interactions using a three-track neural network. Science, 373, 871–876.

49. Abramson, J., Adler, J., Dunger, J., Evans, R., Green, T., Pritzel, A., Ronneberger, O., Willmore, L., Ballard, A.J., Bambrick, J., et al. (2024) Accurate structure prediction of biomolecular interactions with AlphaFold 3. Nature, 10.1038/s41586-024-07487-w.

50. Alfieri, C., Gambetta, M.C., Matos, R., Glatt, S., Sehr, P., Fraterman, S., Wilm, M., Müller, J. and Müller, C.W. (2013) Structural basis for targeting the chromatin repressor Sfmbt to Polycomb response elements. Genes Dev., 27, 2367–2379.

51. Górecki, A., Bonarek, P., Górka, A.K., Figiel, M., Wilamowski, M. and Dziedzicka-Wasylewska, M. (2015) Intrinsic disorder of human Yin Yang 1 protein. Proteins Struct. Funct. Bioinforma., 83, 1284–1296.

52. Ahmad, S., Gromiha, M.M. and Sarai, A. (2004) Analysis and prediction of DNA-binding proteins and their binding residues based on composition, sequence and structural information. Bioinformatics, 20, 477–486.

53. Corley, M., Burns, M.C. and Yeo, G.W. (2020) How RNA-Binding Proteins Interact with RNA: Molecules and Mechanisms. Mol. Cell, 78, 9–29.

54. Ellis, J.J., Broom, M. and Jones, S. (2007) Protein–RNA interactions: Structural analysis and functional classes. Proteins Struct. Funct. Bioinforma., 66, 903–911.

55. Hoffman, M.M., Khrapov, M.A., Cox, J.C., Yao, J., Tong, L. and Ellington, A.D. (2004) AANT: the Amino Acid–Nucleotide Interaction Database. Nucleic Acids Res., 32, D174–D181.

56. Wang, W., Qiao, S., Li, G., Cheng, J., Yang, C., Zhong, C., Stovall, D.B., Shi, J., Teng, C., Li, D., et al. (2022) A histidine cluster determines YY1-compartmentalized coactivators and chromatin elements in phase-separated enhancer clusters. Nucleic Acids Res., 50, 4917–4937.

57. Wang, X., Bigman, L.S., Greenblatt, H.M., Yu, B., Levy, Y. and Iwahara, J. (2023) Negatively charged, intrinsically disordered regions can accelerate target search by DNA-binding proteins. Nucleic Acids Res., 10.1093/nar/gkad045.

58. Holmes, Z.E., Hamilton, D.J., Hwang, T., Parsonnet, N.V., Rinn, J.L., Wuttke, D.S. and Batey, R.T. (2020) The Sox2 transcription factor binds RNA. Nat. Commun., 11.

59. Hansen, A.S., Hsieh, T.S., Cattoglio, C. and Pustova, I. (2018) An RNA-binding region regulates CTCF clustering and chromatin looping.

60. Saldaña-Meyer, R., Rodriguez-Hernaez, J., Escobar, T., Nishana, M., Jácome-López, K., Nora, E.P., Bruneau, B.G., Tsirigos, A., Furlan-Magaril, M., Skok, J., et al. (2019) RNA Interactions Are Essential for CTCF-Mediated Genome Organization. Mol. Cell, 76, 412-422.e5.

61. Hansen, A.S., Hsieh, T.-H.S., Cattoglio, C., Pustova, I., Saldaña-Meyer, R., Reinberg, D., Darzacq, X. and Tjian, R. (2019) Distinct Classes of Chromatin Loops Revealed by Deletion of an RNA-Binding Region in CTCF. Mol. Cell, 76, 395-411.e13.

62. Ficzycz, A. and Ovsenek, N. (2002) The Yin Yang 1 Transcription Factor Associates with Ribonucleoprotein (mRNP) Complexes in the Cytoplasm of Xenopus Oocytes *. 277, 8382–8387.

63. Figiel, M., Szubert, F., Luchinat, E., Bonarek, P., Baranowska, A., Wajda-Nikiel, K., Wilamowski, M., Miłek, P., Dziedzicka-Wasylewska, M., Banci, L., et al. (2023) Zinc controls operator affinity of human transcription factor YY1 by mediating dimerization via its N-terminal region. Biochim. Biophys. Acta BBA - Gene Regul. Mech., 1866, 194905.

64. Xiao, R., Chen, J.Y., Liang, Z., Luo, D., Chen, G., Lu, Z.J., Chen, Y., Zhou, B., Li, H., Du, X., et al. (2019) Pervasive Chromatin-RNA Binding Protein Interactions Enable RNA-Based Regulation of Transcription. Cell, 178, 107-121.e18.

