## Supplemental Figures and Legends for "The N-terminus of YY1 regulates DNA and RNA binding affinity for both the zinc-fingers and an unexpected nucleic acid binding domain"

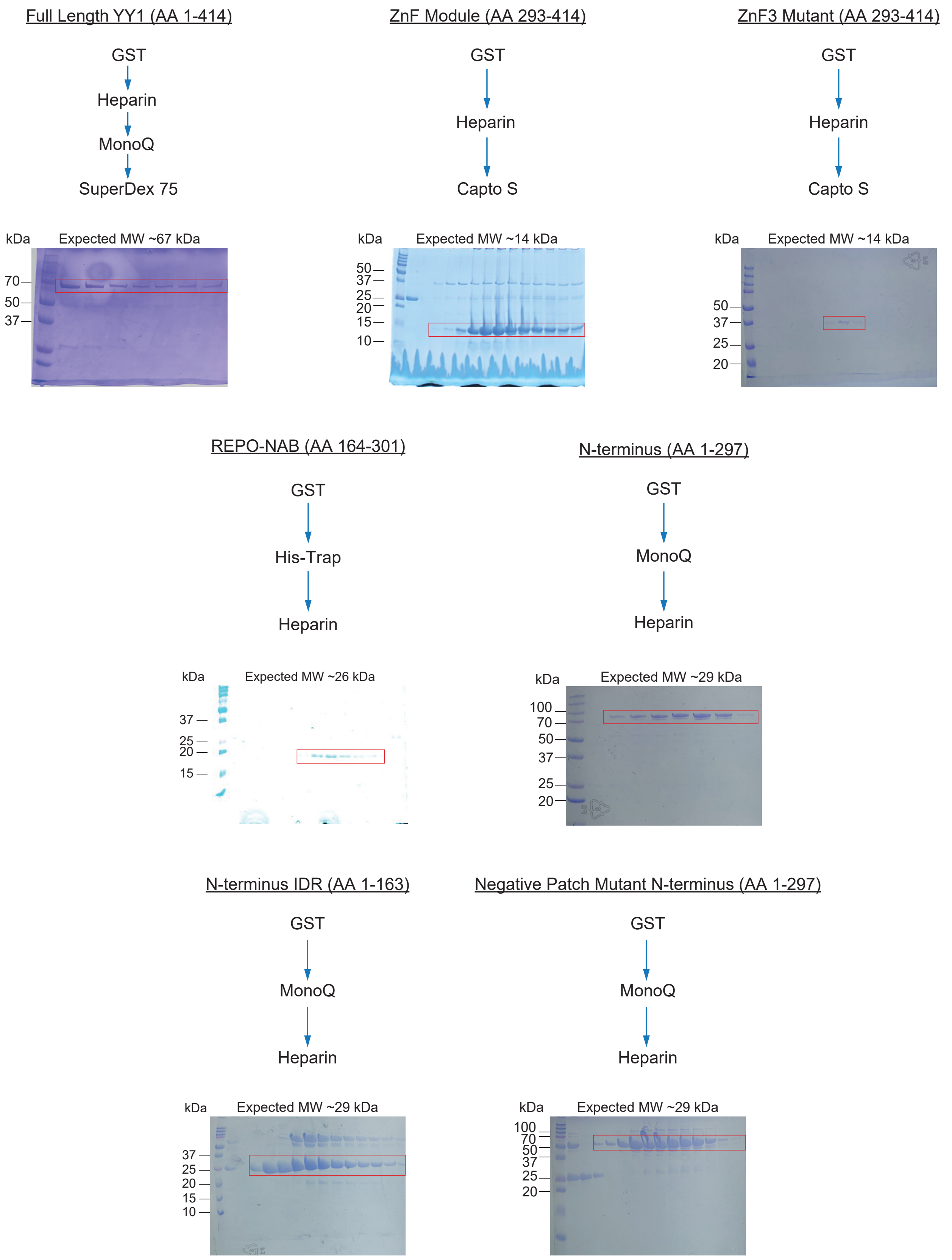


**Figure S1.** Purification schemes for full length YY1 and mutant constructs. Depicted are the step-wise approaches for purification of protein constructs utilized within this paper. Expected MWs (kDa) of each construct is reported above each respective SDS-PAGE gel, which showcase the final purity of the protein purified and utilized in subsequent assays. We note that the N-terminus purification yields a protein that migrates at an elevated MW than anticipated and attribute this deviation to the inherent anomalous SDS-PAGE migration of full length YY1 (16) as well as previously observed peculiar migration of the N-terminus (63).

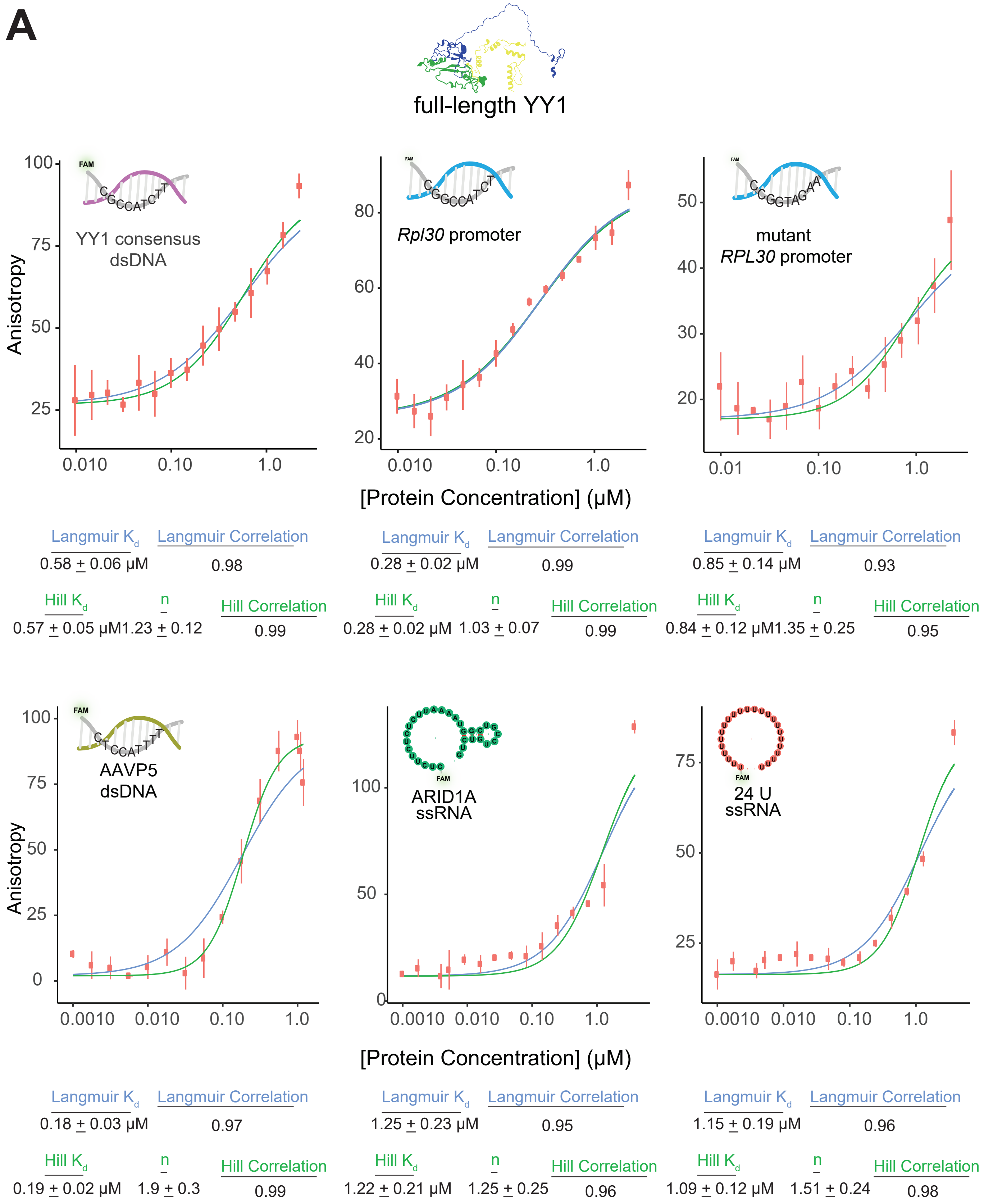


**Figure S2.** Purified full length YY1 exhibits characteristic binding behaviours to our full panel of nucleic acids. **(A)** Fluorescence polarization assays against our panel of nucleic acids. Error bars represent standard deviation from three technical replicates and the nonlinear regression fits of both Langmuir (blue) and Hill (green) equations are depicted for each graph. Langmuir observed $K_d$ measurements are reported below each binding curve, as well as Hill observed $K_d$ and the Hill coefficient (n). Concentration ranges are available within the supplementary datasets uploaded to Zenodo.

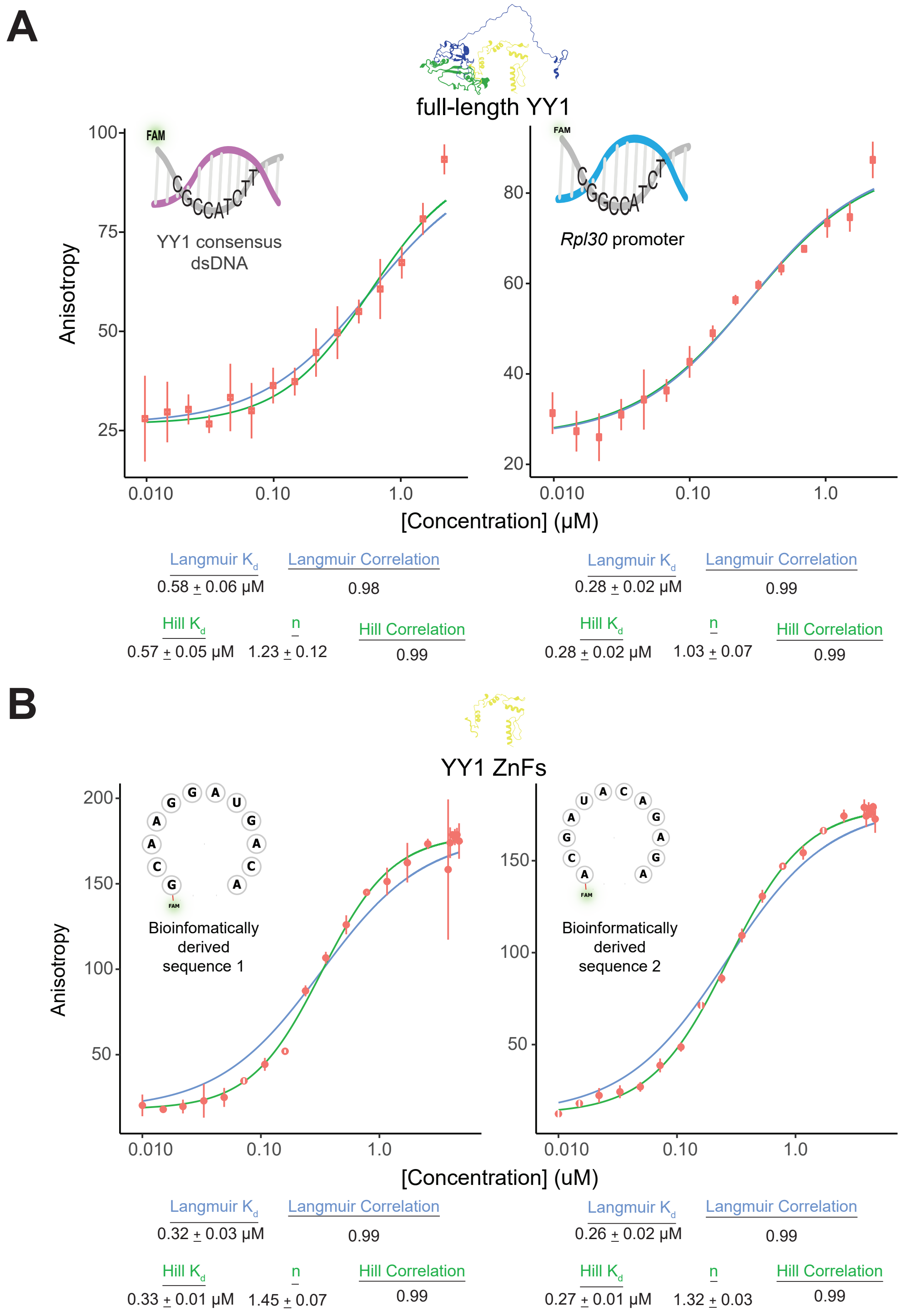


**Figure S3.** Full length YY1 and the ZnF module exhibit characteristic specific activities. **(A)** Full length YY1 fluorescence polarization assays against the YY1 consensus dsDNA sequence and the *Rpl30* dsDNA promoter. Error bars represent standard deviation from three technical replicates and the nonlinear regression fits of both Langmuir (blue) and Hill (green) equations are depicted for each graph. Langmuir observed $K_d$ measurements are reported below each binding curve, as well as Hill observed $K_d$ and the Hill coefficient (n). Full length YY1 concentration range: 10 nM – 1.5 μM. **(B)** Fluorescence polarization assays utilizing the ZnFs against two bioinformatically derived ssRNA sequences from Sigova et al. (27) CLIP-seq data. The methodology to identify these sequences is outlined in the Methods section “Computational analyses of YY1 genomic binding sites”. Error bars represent standard deviation from three technical replicates and the nonlinear regression fits of both Langmuir (blue) and Hill (green) equations are depicted for each graph. Langmuir observed $K_d$ measurements are reported below each binding curve, as well as Hill observed $K_d$ and the Hill coefficient (n).

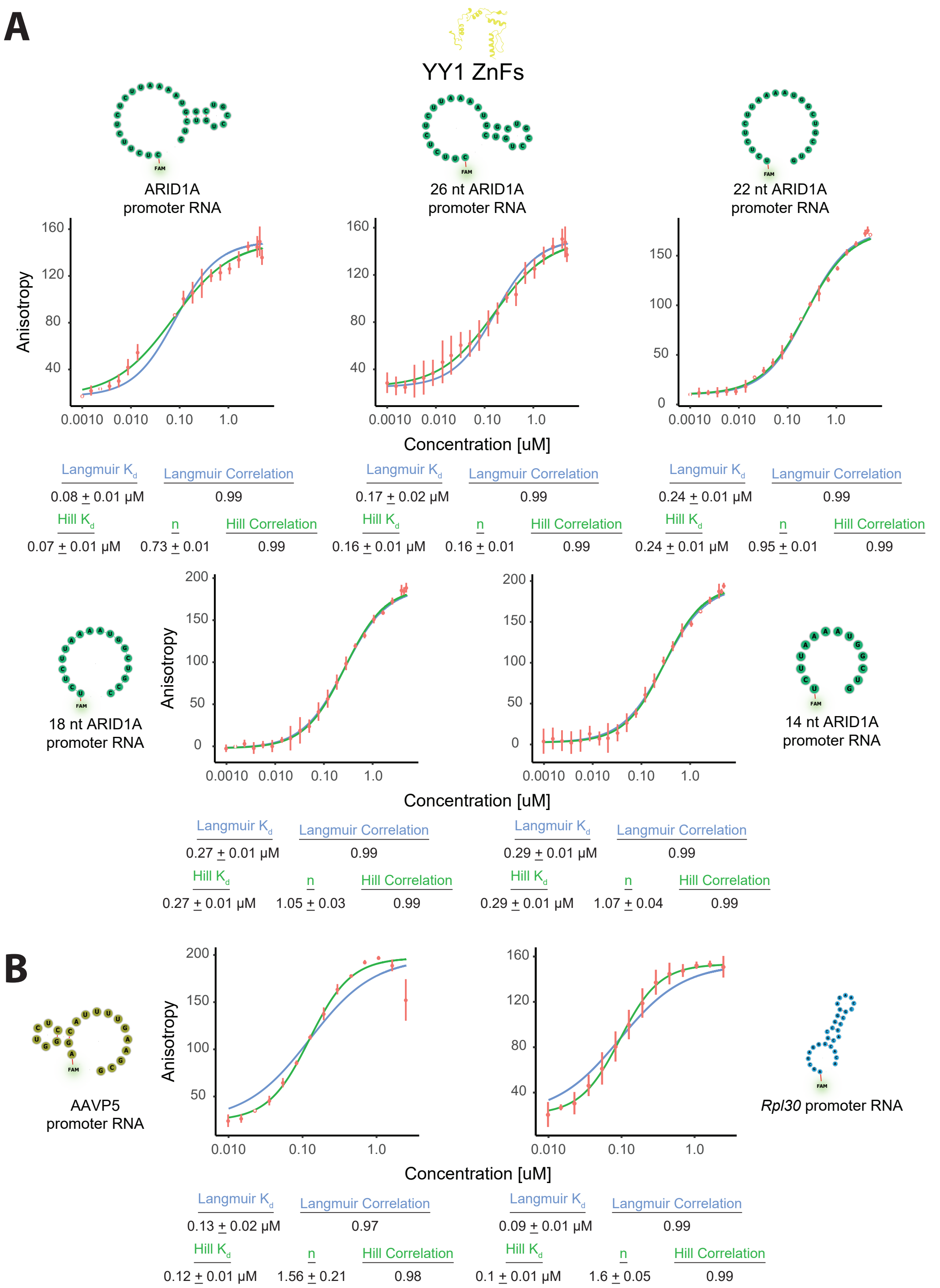


**Figure S4.** YY1's ZnF module displays little sequence, length, or secondary structure specificity in binding ssRNA. **(A)** Fluorescence polarization binding curves utilizing YY1's ZnF module and a truncation series of *ARID1A* ssRNAs. The initial *ARID1A* ssRNA (top left) is the 30-nt nucleic acid substrate consistently used within our work, and subsequent truncations remove 2 nucleotides from both the 5' and 3' ends of the previous ssRNA, rendering shorter ssRNA molecules of lengths 26, 22, 18, and 14 nt. Error bars represent standard deviation from three technical replicates and the nonlinear regression fits of both Langmuir (blue) and Hill (green) equations are depicted for each graph. Langmuir observed $K_d$ measurements are reported below each binding curve, as well as Hill observed $K_d$ and the Hill coefficient (n). **(B)** Fluorescence polarization binding curves utilizing YY1's ZnF module and ssRNA species derived from their respective dsDNA substrates i.e. these ssRNAs are the transcriptional products of the AAVP5 dsDNA promoter and the *Rpl30* dsDNA promoter. Error bars represent standard deviation from three technical replicates and the nonlinear regression fits of both Langmuir (blue) and Hill (green) equations are depicted for each graph. Langmuir observed $K_d$ measurements are reported below each binding curve, as well as Hill observed $K_d$ and the Hill coefficient (n).

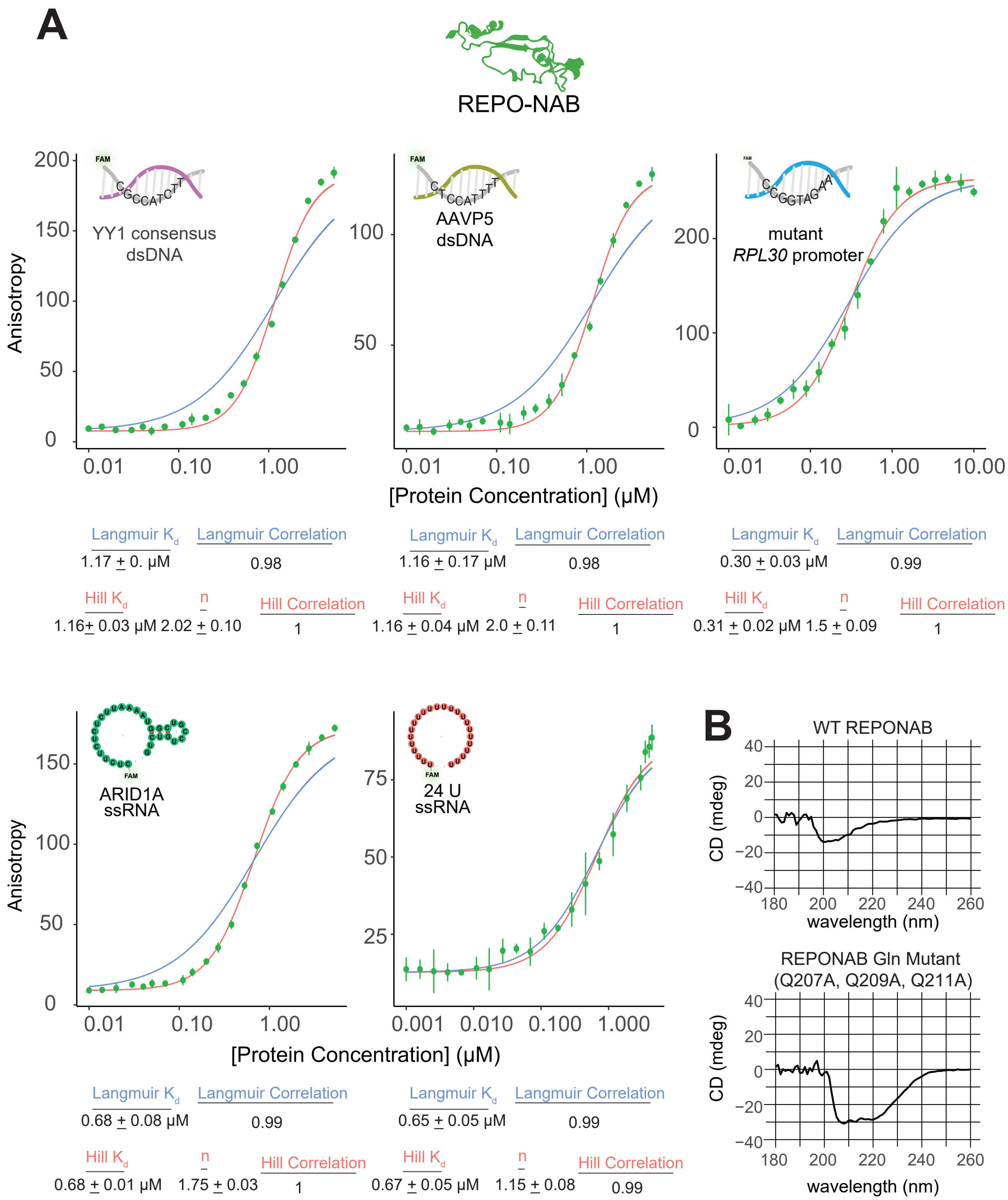


**Figure S5.** The REPO-NAB domain binds nucleic acids and maintains its β-sheet character in isolation. **(A)** Fluorescence polarization assays against our panel of nucleic acids. Error bars represent standard deviation from three technical replicates and the nonlinear regression fits of both Langmuir (blue) and Hill (red) equations are depicted for each graph. Langmuir observed $K_d$ measurements are reported below each binding curve, as well as Hill observed $K_d$ and the Hill coefficient (n). Concentration ranges are available within the supplementary datasets uploaded to Zenodo. **(B)** CD spectroscopy of WT REPO-NAB and the 3Q→A mutant demonstrating both adopt similar β-sheet rich folds at room temperature. Note that the concentration of the 3Q→A mutant is somewhat higher.

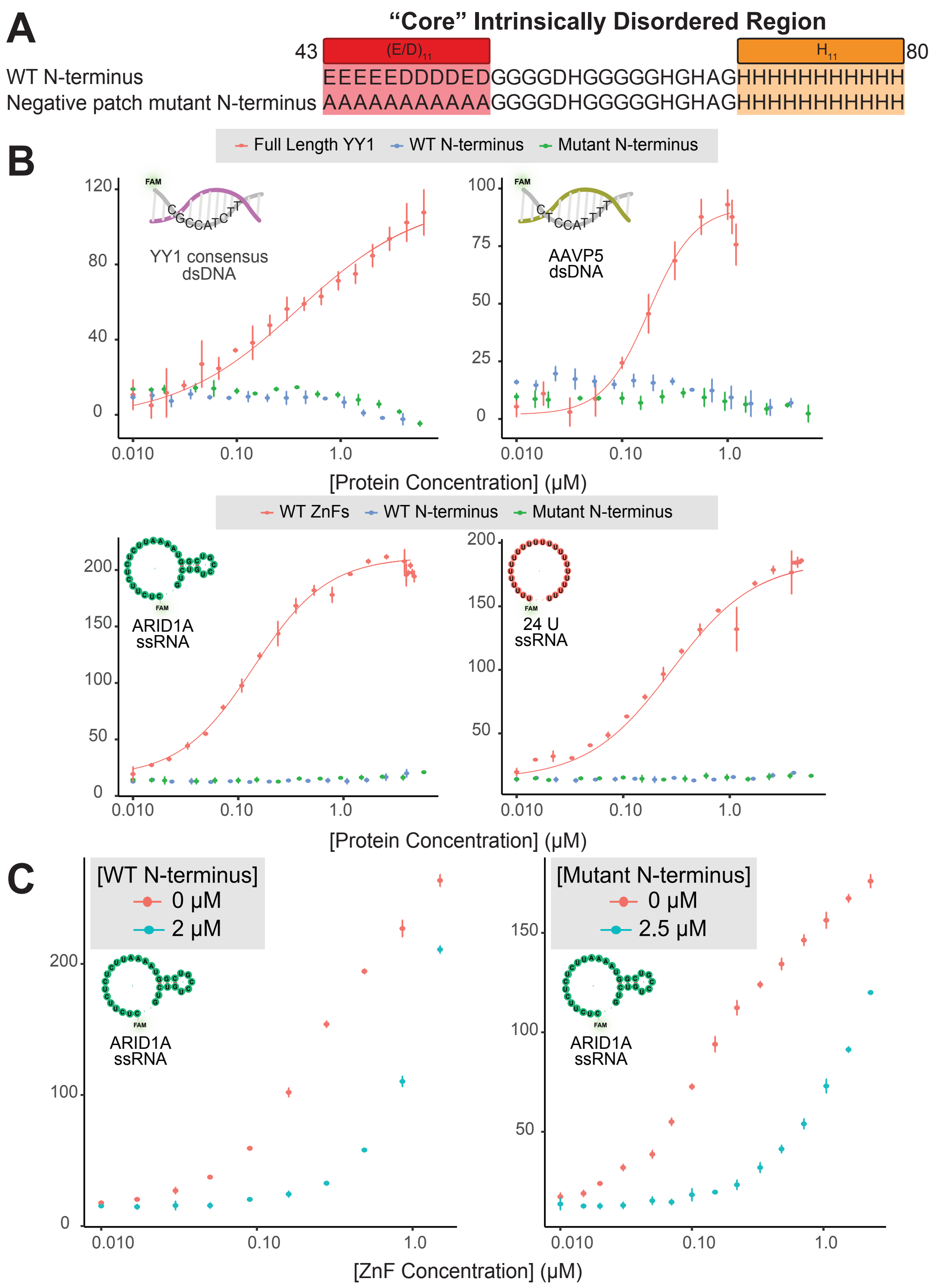


**Figure S6.** The acidic stretch (AA 43-53) of YY1's N-terminus is dispensable for N-terminus mediated autoregulation of ZnF nucleic acid binding. **(A)** Single letter amino acid representation of both the wildtype (WT) and negative patch mutant of the "core Intrinsically Disordered Region (IDR)" (56) in YY1's N-terminus. **(B)** Fluorescence polarization binding experiments for the indicated protein constructs and nucleic acids. Error bars represent standard deviation across three technical replicates. **(C)** Fluorescence polarization competition experiments. (Left) Competition experiment from figure 5A, highlighting the highest concentration of WT N-terminus inhibiting ZnF nucleic acid binding. (Right) The indicated amounts of purified negative stretch mutant N-terminus were added in *trans* to purified ZnFs and binding to the representative nucleic acid substrates was measured. Error bars represent standard deviation across three technical replicates.

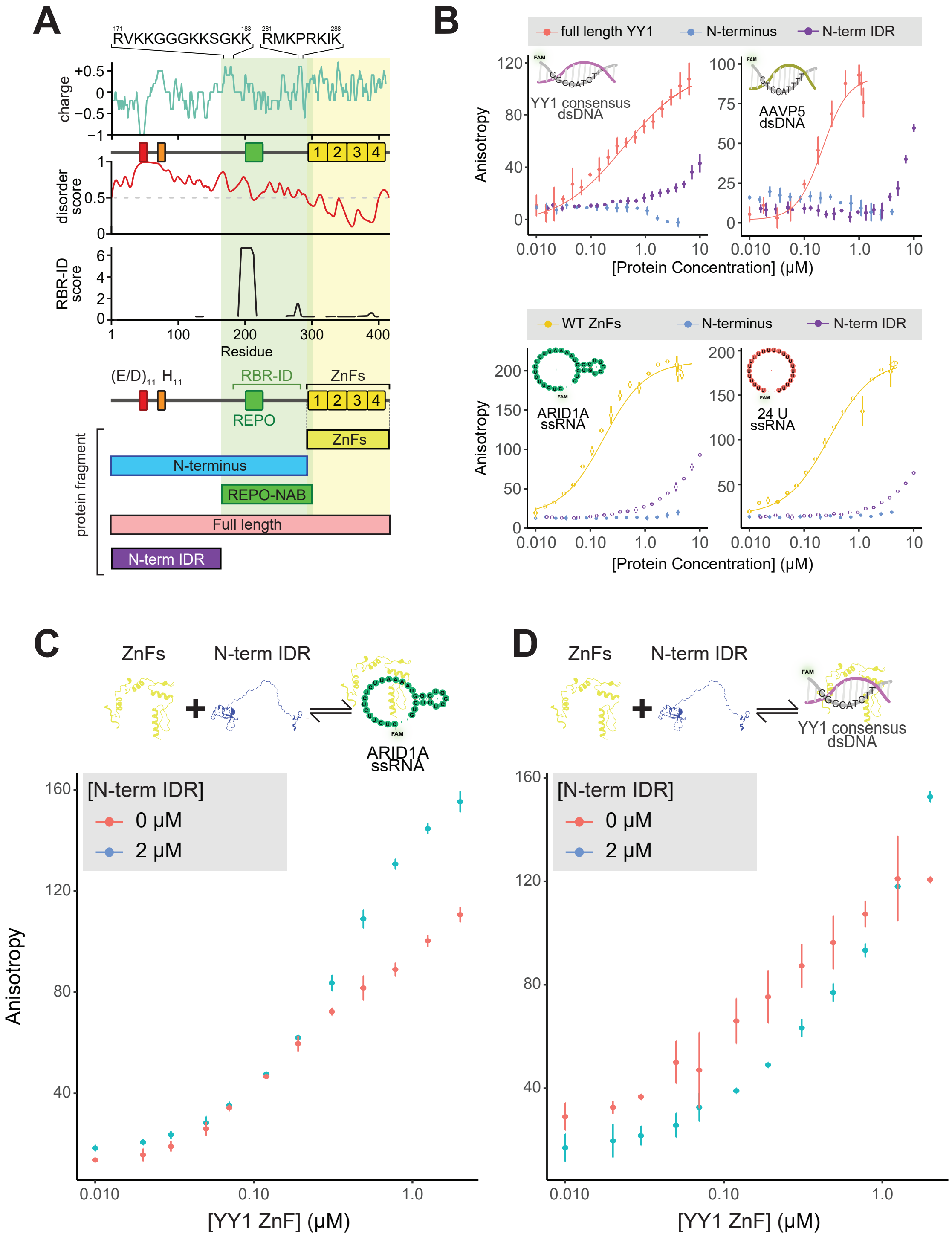


**Figure S7.** YY1's N-terminal IDR (N-terminus ΔREPO) has little nucleic acid binding capacity and does not inhibit ZnF nucleic acid binding. **(A)** (Top) YY1 domain map. EMBOSS net charge for 5 amino acid sliding window, with the only two regions highlighted that are basic patches not in the ZnF module as preciously defined (5 or more consecutive resides that have K/R represented at a frequency >0.5, also termed "ARM-like" motifs (28). (Upper Middle) IUPRED3 disorder prediction spanning YY1. (Lower Middle) RNA Binding Region (RBR)-ID score (46) displaying the amino acid residues significantly cross-linked to nuclear RNA within E14 mESCs. (Bottom) Domain structure and pertinent protein constructs that have been purified and assayed, note that the N-terminus span (AA 1-297) is the same as in Sigova et al. **(B)** Fluorescence polarization binding experiments for the indicated protein constructs and nucleic acids. Error bars represent standard deviation across three technical replicates. **(C)** and **(D)** Fluorescence polarization competition experiments. The indicated amounts of purified ΔREPO N-terminus were added in *trans* to purified ZnFs and binding to the representative nucleic acid substrates was measured. Error bars represent standard deviation across three technical replicates.
